# The ovarian cycle and estradiol modulate cortical network state and auditory entrainment in mice and women

**DOI:** 10.64898/2026.10.12.757951

**Authors:** Maria Isabel Carreño-Muñoz, Pegah Chehrazi, Alessandra Ciancone Chama, Inga Knoth, Antônia Samia Fernandes do Nascimento, Benjamin F.N. Campbell, Jozsef Csicsvari, Bidisha Chattopadhyaya, Sarah Lippé, Graziella Di Cristo

## Abstract

The human brain is remarkably dynamic, continuously adapting to internal and external signals across rapid and protracted timescales. While both male and female brains are influenced by sex hormones to drive adaptive behavioural and physiological changes, the ovarian cycle in female mammals represents a conserved naturalistic model to understand how hormones such as estradiol (E2) alter cellular and network states to shape brain function.

To understand whether and how the ovarian cycle influences sensory cortical processing, we investigated cortical network activity at rest and upon auditory stimulation across the ovarian cycle in mice and human subjects. We recorded EEG activity in men and women and intracortical local field potentials in male and female mice. Female participants were tested during two phases of their ovarian cycle characterized by low E2 levels (early follicular phase in women; late estrus in mice) and high E2 levels (late follicular phase in women; proestrus in mice).

Across species, high endogenous or exogenous estradiol was associated with a steeper aperiodic power spectral slope, indicating decreased excitatory/inhibitory drive and attenuated auditory entrainment following a 40Hz stimulation. At the single-cell level, high E2 selectively increased the baseline and sustained 40-Hz firing of putative fast-spiking interneurons while reducing their temporal coherence during both spontaneous and stimulus-evoked activity, suggesting that E2 enhances inhibitory tone, while disrupting the temporal coordination of inhibitory networks. Altogether, these findings demonstrate that estradiol dynamically modulates cortical network states and auditory responsiveness. This underscores the necessity of investigating hormone-brain coupling as an evolutionarily conserved model essential for yielding translational insights into sex-specific neurobiology.

## INTRODUCTION

Internal physiological states, such as those determined by endocrine rhythms, play a crucial role in modulating neural circuits in the brain, thereby shaping sensory perception, sensorimotor coordination, and learning^1^. In mammals, circulating levels of ovarian sex steroid hormones fluctuate throughout the ovarian cycle, thereby intricately orchestrating physiological states and behaviors^2–4^. While extensive research has been devoted to understanding how circulating sex hormones affect cognitive function and behavior^5^, their impact on neuronal activity and network states is poorly understood precluding true mechanistic insight.

Sex steroid hormones such as 17β-oestradiol (E2), the most bioactive form of estrogens in non-pregnant mammals, are recognized as key neuromodulators shaping brain organisation and function. In fact, since seminal studies three decades ago demonstrated that E2 regulates dendritic spine density on CA1 pyramidal neurons in the hippocampus^6^ and potentiates hippocampal excitatory synaptic plasticity^7,8^, an extensive body evidence has firmly established the importance of E2 as a critical modulator of learning and memory^5,9^. Beyond the hippocampus, E2 can modulate neuronal excitability and synaptic function in sensory cortices in a neuron-type and dose-dependent manner^10–13^. Yet, how physiological E2 fluctuations shape network states and influence responses to sensory stimuli at cellular and population levels remains unclear. Furthermore, whether this hormone-dependent modulation represents an evolutionarily conserved principle of neural function across species is unknown.

Here, we investigated the effects of the ovarian cycle on cortical network activity at rest and as evoked by auditory stimulation, using longitudinal recording of electroencephalograms (EEGs) in men and women and intracortical local field potentials in male and female mice. Women and female mice were recorded at different phases of the ovarian cycle. In both species, high E2 cycle phases were associated with a steeper aperiodic exponent, reflecting a shifting of resting state network activity toward higher ratio of low vs high frequency power, together with a weaker auditory entrainment to 40-Hz steady-state stimulation. In mice, exogenous E2 administration rapidly increased the steepness of the aperiodic spectral slope and attenuated auditory-evoked entrainment specifically in females, but not males. We further investigated the effects of estrous cycle and estradiol on single-unit firing in mice. We observed estrous phase–dependent changes in putative fast spiking inhibitory neurons; specifically, when endogenous E2 levels were high, these neurons exhibited higher firing rates alongside reduced temporal precision, both at rest and during 40 Hz auditory stimulation. Further, we showed that exogenous E2 administration rapidly and significantly reduced their temporal precision. Altogether, these findings demonstrate that estradiol dynamically modulates cortical network states and auditory responsiveness in a sex-dependent manner, highlighting an evolutionary conserved role for E2 in auditory cortical processing across rodents and primates.

## RESULTS

### Estradiol levels shape cortical resting-state spectral dynamics across species

Brain activity at rest is shaped by the ongoing balance of excitation and inhibition within cortical circuits in the absence of stimulus presentation. Several studies reported that estradiol and other ovarian hormone levels modulate gamma-band activity during resting state, passive sensory stimulation and cognitive tasks in both rodents^14–20^ and humans^21–25^. However, their impact on the spectral organization of spontaneous cortical activity and to what extent this effect is evolutionary conserved in mammals remains poorly understood, due to contrasting results ^14–25^. This confounding variability could be due to differences in experimental design (e.g., one vs multiple longitudinal recordings), stimulation type and protocols, data processing and analysis pipeline.

To clarify the impact of the ovarian cycle on resting state and sensory evoked cortical network activity, we conducted longitudinal recordings across the reproductive cycle in female mice and women, alongside repeated recordings at matched time intervals in male participants and mice. First, we quantified resting state cortical activity in both species. In humans, we recorded EEG across the scalp using a 128-channel system; for the analyses presented here, we focused on Cz and the surrounding electrodes, a central cluster selected to capture auditory cortical responses^26,27^ In women, recordings were acquired consecutively at two defined menstrual cycle phases: self-reported cycle day 1 (Figure 1A; EF, early follicular phase, low E2) and 10 days later (Figure 1A; LF, late follicular phase, high E2). Men were recorded twice, 10 days apart, at the same time of the day. In mice, we recorded intracortical LFP from the primary auditory cortex (A1) using chronically implanted tetrodes. Females were recorded multiple times across estrous phase for at least 3 complete estrous cycles (Figure 1B; Supplemental Figure 1, 2). Males were recorded at comparable intervals (daily or every other day) during 3 weeks; for the analyses presented here, two recording sessions separated by two days in the same familiarized environment were selected to provide a time-matched control for repeated recordings. Human resting state activity was acquired under controlled open-eye conditions during a visual cross-fixation task, in the absence of auditory input. In mice, locomotor activity varied across the estrous cycle, with females exhibiting increased locomotion during proestrus compared to estrus (Supplementary Figure 3). Therefore, to minimize locomotion- and state-dependent variability and better approximate human resting conditions, analyses were restricted to spontaneous immobility epochs (body center displacement < 2 cm/s for more than 2 seconds).

**Figure 1.**
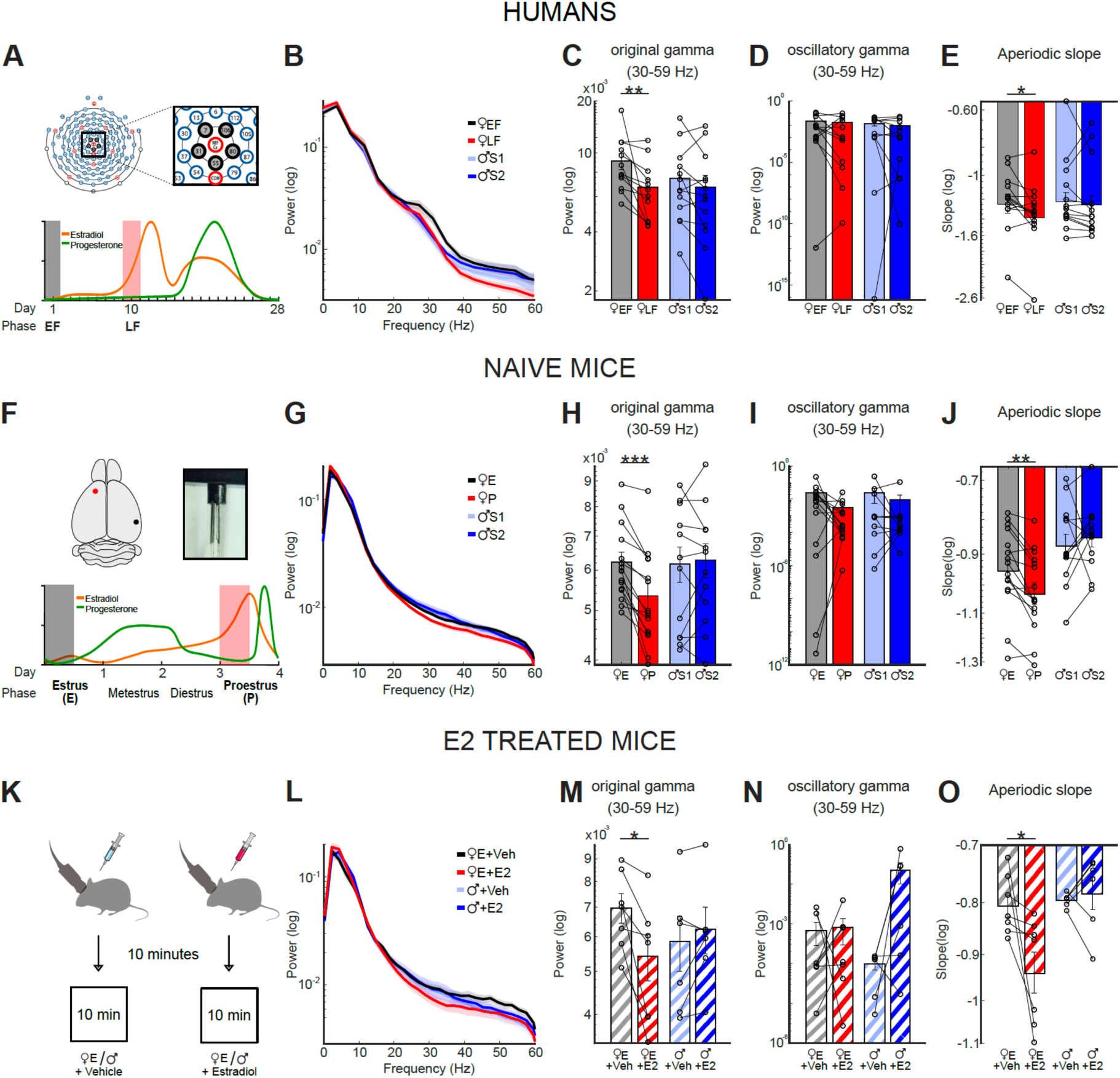
Higher estradiol levels are associated with lower aperiodic baseline gamma power across species. [A] Experimental design in humans. Top: Electrode configuration (chosen recording electrodes in black, reference in red). Bottom: Schematic of estradiol and progesterone fluctuations across the ovarian cycle in humans, indicating low (black shadow) and high (red shadow) E2 phases. [B] Classical power spectral density (PSD; 0–60 Hz) computed from 4-min resting-state EEG recordings in humans. The logarithmic scale is used for visualization purposes only. [C] Low-gamma bands in humans. women were compared between low-E2 (early follicular, EF) and high-E2 (late follicular, LF) phases, and men were compared between two recording sessions separated by 10 days (S1 vs S2). Wilcoxon signed-rank tests for paired samples: women EF vs LF: z = 3.1798, p **=** 0.0015; men S1 vs S2: z = 0.9435, p = 0.3454. [D] Periodic low-gamma component following FOOOF decomposition of the power spectra in humans Wilcoxon signed-rank tests for paired samples: women EF vs LF: z = 0.7060, p = 0.4802; men S1 vs S2: z = 0.2446, p = 0. 8068). [E] Aperiodic slope component following FOOOF decomposition of the power spectra in humans. Wilcoxon signed-rank tests for paired samples: women EF vs LF: z = 2.2014, p = 0.0277; men S1 vs S2: z = 1.2929, p = 0. 1961. [F] Experimental design in mice. Top: Electrode configuration (chosen recording electrodes in black, reference in red). Bottom: Schematic of estradiol and progesterone fluctuations across the ovarian cycle in humans, indicating low (black shadow) and high (red shadow) E2 phases. [G] Classical power spectral density (PSD; 0–60 Hz) computed from immobility periods (>2 s) during 10 min of recordings in freely behaving mice. [H] Low-gamma bands in mice. Females in estrus (low E2) were compared to females in proestrus (high E2), and males were compared between two recording sessions in an already familiarized open field (S1 vs S2). Wilcoxon signed-rank tests for paired samples: females in proestrus vs estrus: z=3.4128 p=0.0006; males S1 vs S2: z=-0.3922 p=0.6949. [I] Periodic low-gamma component following FOOOF decomposition of the power spectra in naïve mice. Wilcoxon signed-rank tests for paired samples: females in proestrus vs estrus: z=1.5335 p=0.1252; males S1 vs S2: z=-0.3922 p=0.6949. [J] Aperiodic slope component following FOOOF decomposition of the power spectra in naïve mice. Wilcoxon signed-rank tests for paired samples: females in proestrus vs estrus: z=3.0670 p=0.0021; males S1 vs S2: z=-0.7844 p=0.4327. [K] Experimental scheme of pharmacological approach. Females in proestrus (high E2) and male mice were intraperitoneally injected with either vehicle or E2 (20 µg/kg, i.p.), left rest in their home cage for 10 minutes and then recorded for another 10 minutes of spontaneous behaviour. [L] Classical power spectral density (PSD; 0–60 Hz) computed from immobility periods (>2 s) during 10 min of recordings in freely behaving mice. [M] Low-gamma bands in treated mice. After recordings at different estrous cycle phases, male mice and female mice in estrus were injected with vehicle and E2 (in randomized order). Wilcoxon signed-rank tests for paired samples: females in estrus+Veh vs females in estrus+E2: z=2.3664 p=0.0180; males+Veh vs males+E2 z=-0.9435 p=0.3454. [N] Periodic low-gamma component following FOOOF decomposition of the power spectra in E2 treated mice. Wilcoxon signed-rank tests for paired samples: females in estrus+Veh vs females in estrus+E2: z=-0.5071 p=0.6121; males+Veh vs males+E2 z=-1.7821 p=0.0747. [O] Aperiodic slope component following FOOOF decomposition of the power spectra in E2 treated mice. Wilcoxon signed-rank tests for paired samples: females in estrus + Veh vs females in estrus + E2: z=2.3664 p=0.0179; males +Veh vs males+E2 z=-0.3145 p=0.7532. *P < 0.05. **P < 0.01. EF: early follicular; LF: late follicular; E: estrous; P: proestrus. Bar plots show mean ± SEM. Dots represent individual data. Number of human participants: women = 13, men = 13; number of naïve mice: females = 16, males =12; number of treated mice: females = 7, males =6.

Conventional power spectral density (PSD) analysis revealed significantly lower resting state gamma band power during higher-E2 states, in both women (Figure 1D,G; late follicular phase) and female mice (Figure 1E, H; proestrus) compared with their respective lower-E2 conditions (early follicular phase in women and estrus in mice). Conversely, no significant differences were observed across repeated recording sessions in men or male mice (Figure 1D,E,G,H), indicating that recording order or repeated exposure to the experimental conditions did not affect oscillation power. We further observed that the delta-band power was significantly increased exclusively in female mice at proestrus compared to estrus, while no changes were detected in theta or alpha bands across species (Supplementary Fig. 4). Conventional frequency-band power reflects the combination of oscillatory activity and the broadband aperiodic component of the power spectrum, which can reflect distinct features of underlying cortical network dynamics. We therefore asked whether the ovarian-cycle-dependent changes observed in broadband gamma power reflected modulation of genuine oscillatory activity, changes in the underlying aperiodic component, or a combination of both. To address this point, we next parameterized the spectra using Fitting Oscillations & One-Over-F (FOOOF)^28^, which separates periodic oscillatory activity from the underlying aperiodic 1/f component. After subtracting the fitted aperiodic component from the power spectrum, gamma oscillatory power did not significantly differ between low- and high-E2 conditions (Figure 1 J, K). In contrast, we observed a consistent estrous cycle-dependent change in the aperiodic component of the spectrum, with higher-E2 states characterized by a steeper 1/f spectral slope indicating a higher relative distribution of power in low versus high frequencies (Figure 1 M, N). No comparable session-dependent modulation was observed in men or male mice (Figure 1 M, N). Thus, the apparent decrease in gamma-band power measured from the conventional PSD during high-E2 states primarily reflected a broadband reorganization driven by the aperiodic component of the power spectrum rather than a selective reduction in gamma oscillatory activity.

To directly test whether estradiol was sufficient to induce this spectral reorganization, we administered acute systemic E2 (20 µg/kg, i.p.) or vehicle (saline) 15 min before resting-state recordings, during late estrus, when endogenous E2 levels are low (Figure 1C). E2 administration reproduced the endogenous cycle-dependent phenotype in female mice, decreasing gamma oscillation power (Figure 1F, I) and increasing the steepness of aperiodic spectral slope relative to vehicle treatment, while leaving the oscillatory gamma component largely unchanged (Figure 1 L, O). In males, E2 did not reproduce this spectral phenotype (Figure 1F, I, O).; instead, we observed a strong trend toward an increase in the periodic gamma component relative to vehicle treatment (p = 0.0747; Figure 1L).

All together, these results identify the aperiodic component, rather than gamma oscillations per se, as a robust spectral signature of estradiol-dependent modulation of baseline cortical state in mice, with a similar pattern observed in humans during low estradiol associated cycle phases.

### Estradiol levels affect the temporal organisation of narrow-spiking neuron firing

Our data show that higher E2 levels are associated with a steeper aperiodic spectral slope, a network signature previously linked to a shift toward stronger inhibitory tone^28–30^. To test whether estrous cycle-dependent changes in population activity were accompanied by distinct alterations in single-cell spontaneous firing across different neuronal subtypes, we performed extracellular tetrode recordings in A1 layer 5 of adult cycling female mice (Figure 2A). We classified single units based on waveform width into broad-spiking (putative excitatory neurons) and narrow-spiking (putative fast-spiking inhibitory interneurons)^31^(Figure 2B). When considering the full population, baseline firing rates did not vary across the estrous cycle (Figure 2C). However, separating neurons by subtype revealed cell type–specific effects: broad-spiking (BS) neurons showed no modulation across phases, whereas narrow-spiking (NS) neurons exhibited a significant increase in firing rate during proestrus (Figure 2 C). This selective increase in NS neurons activity during the high-E2 phase is consistent with the steeper aperiodic slope observed in the FOOOF decomposition of the LFP power spectra, thus supporting the hypothesis that inhibitory activity is higher during proestrus than estrus.

**Figure 2.**
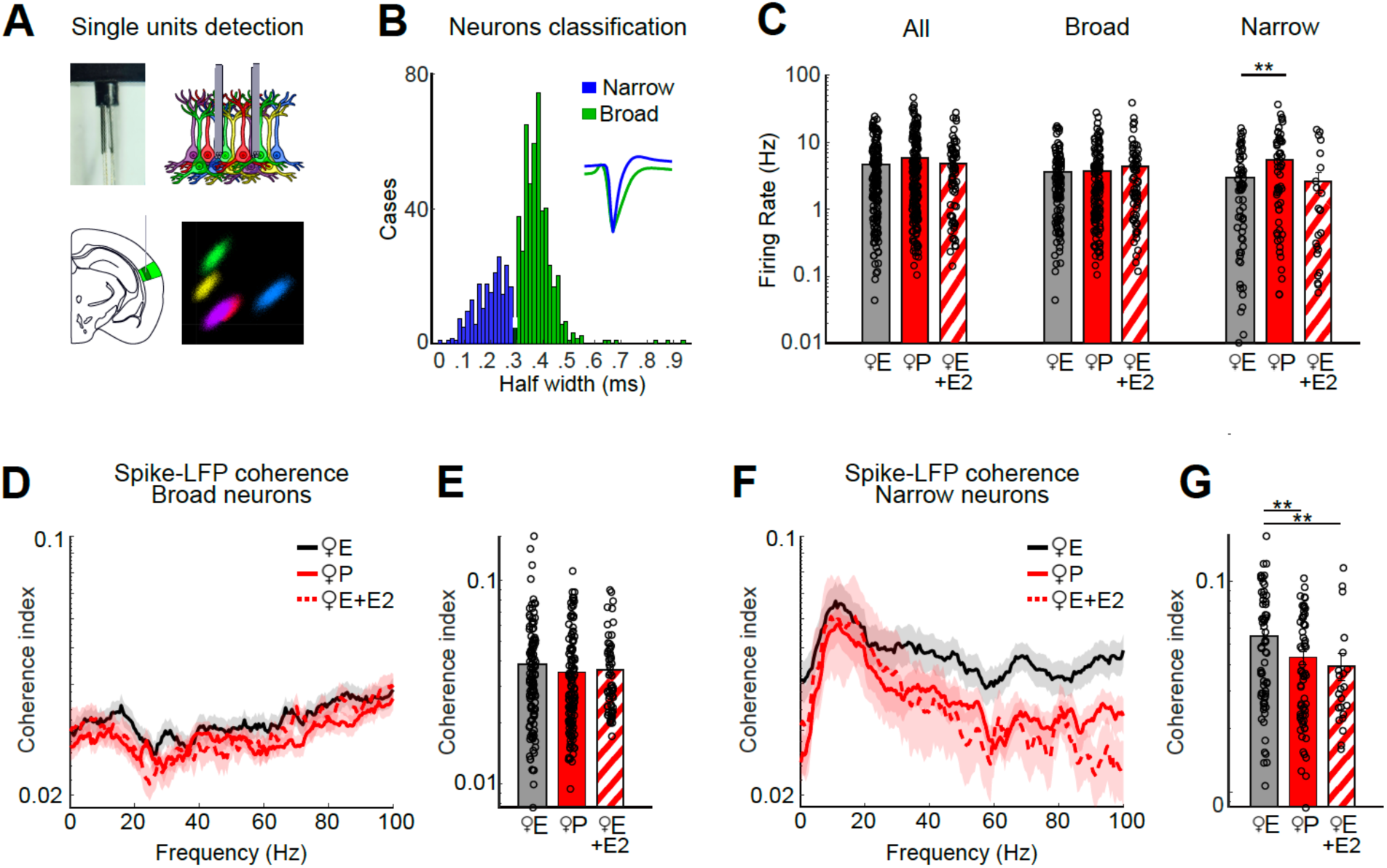
High estradiol phase is associated with higher firing rates but reduced spike–LFP coherence in narrow-spiking neurons in mice. [A] Single-unit recording experimental approach. Top left: Photograph of a microdrive containing four tetrodes. Top right: Schematic illustrating tetrode recording surrounding neurons. Bottom left: Schematic of the targeted cortical region (primary auditory cortex, A1). Bottom right: Example of spike-sorting and clustering analysis used for single-unit identification. [B] Distribution of spike waveform widths across all recorded single units. The bimodal distribution was used to classify neurons into broad (green) vs narrow (blue) spiking populations. Insets show mean normalized spike waveforms for each population. [C] Mean firing rate during immobility periods (>2 s) within the first 10 min of exposure to a familiar open field. Statistical analyses were performed using linear mixed-effects models (LMM) to account for variability across cells and animals. Comparisons are shown for all neurons (estrus vs proestrus: p = 0.1401; estrus vs estrus+E2: p = 0.8684), broad neurons (estrus vs proestrus: p = 0.9731; estrus vs estrus+E2: p = 0.2299) and narrow neurons (estrus vs proestrus: p = 0.0084; estrus vs estrus+E2: p = 0.7514). [D] Spike–LFP coherence spectrograms for broad-spiking neurons. [E] Mean spike–LFP coherence in the broadband gamma range (30–100 Hz) for broad-spiking neurons. LMM, estrus vs proestrus: p = 0.7120; estrus vs estrus+E2: p=0.6001. [F] Spike–LFP coherence spectrograms for broad-spiking narrow-spiking neurons. [G] Mean spike–LFP coherence in the broadband gamma range (30–100 Hz) for narrow-spiking neurons. LMM, estrus vs proestrus: p = 0.0022; estrus vs estrus+E2: p=0.0081. **P < 0.01. E: estrous; P: proestrus. Bar plots show mean ± SEM. Dots represent individual neuron data. Number of female mice = 13. All neurons N: estrus = 178, proestrus= 195, estrus + E2 = 94. Broad Spiking neurons number: estrus = 110, proestrus= 136, estrus + E2 = 70. Narrow Spiking cell number: estrus= 68, proestrus= 59, estrus+E2= 24. N female mice=13.

Despite this elevated firing activity, spike–LFP coherence analyses uncovered a dissociation between firing output and network coupling. Specifically, NS neurons displayed reduced spike-LFP coherence, particularly in the broadband gamma range, during proestrus compared to estrus, while BS neurons again showed no cycle-dependent changes (Figure 2 D-G). Thus, although NS neurons were more active during proestrus, their spikes were less consistently aligned with ongoing local network oscillations. Reduced spike–LFP coherence could reflect a reorganization of the temporal structure of neuronal spiking, changes in the population-level dynamics captured by the LFP, or both, ultimately resulting in a less consistent temporal relationship between single-neuron firing and ongoing network activity.

Next, we examined whether acute E2 administration was sufficient to promote singe-cell spontaneous firing changes. We found that this treatment specifically reduced spike–LFP coherence of NS neurons, resembling the temporal reorganization observed during proestrus (Figure 2D-G), whereas no significant changes were observed in vehicle-treated mice (Supplementary Fig. 5).

Together, these findings suggest that physiological high-E2 states are characterized by both increased recruitment of NS neurons, which are thought to be putative fast-spiking inhibitory neurons, and reduced temporal coupling of their activity to ongoing network dynamics, whereas acute E2 is sufficient to rapidly reorganize spike–network coupling without altering overall neuronal firing rates.

### Higher estradiol reduces the temporal precision of auditory responses evoked by 40Hz stimulation across species

The hormone-dependent tuning of spontaneous cortical activity we observed likely defines the physiological regime within which sensory information is processed, raising the question of how such changes influence the encoding of incoming sensory inputs. We thus next asked whether and how the ovarian cycle modulates the temporal encoding of incoming sensory information. Auditory steady-state responses (ASSRs) provide a well-established measure of neural entrainment which reflects the brain’s ability to track regularities in rhythmic auditory input and synchronize ongoing neural activity to an external periodic stimulus^32,33^. We therefore used 10 and 40-Hz auditory stimulation to determine whether E2-dependent changes in intrinsic temporal organization influence the capacity of cortical networks to consistently align their activity with an incoming auditory rhythm (Figure 3A). Time–frequency spectrograms revealed that auditory steady-state stimulation at 40 Hz elicited robust oscillatory activity at the stimulation frequency in both humans and mice, with comparable response magnitudes across ovarian cycle stages (Supplementary Fig. 6). However, despite showing similar evoked power, analysis of inter-trial coherence revealed that the temporal precision of these responses across trials was modulated by ovarian stage. Specifically, women in the early follicular phase and female mice in estrus, both characterized by lower endogenous E2 levels, exhibited significantly stronger phase locking to the 40-Hz stimulus than during the corresponding higher-E2 states (late follicular phase in women and proestrus in mice; Fig. 3 B-G), as quantified by inter-trial coherence (ITC) of EEG and LFP signals, respectively. Thus, elevated E2 was associated not with a weaker oscillatory response, but with reduced trial-to-trial temporal precision in the phase alignment of cortical activity to the rhythmic stimulus. On the other hand, men and male mice did not show significant differences in their auditory responses across different recording sessions (Figure 3B-G). In addition, acute E2 administration (20 µg/kg, i.p.) delivered to female, mice during late estrus 25 minutes before the auditory steady-state response paradigm, significantly reduced 40-Hz ITC compared with vehicle-treated controls (Figure 3 H-J), recapitulating the lower phase locking observed during naturally occurring high-E2 states. In contrast, acute E2 administration did not significantly affect auditory ITC in male mice (Figure 3H-J). Furthermore, entrainment to slower 10-Hz stimulations was unaffected by estrous cycle (Supplementary Fig. 7).

**Figure 3.**
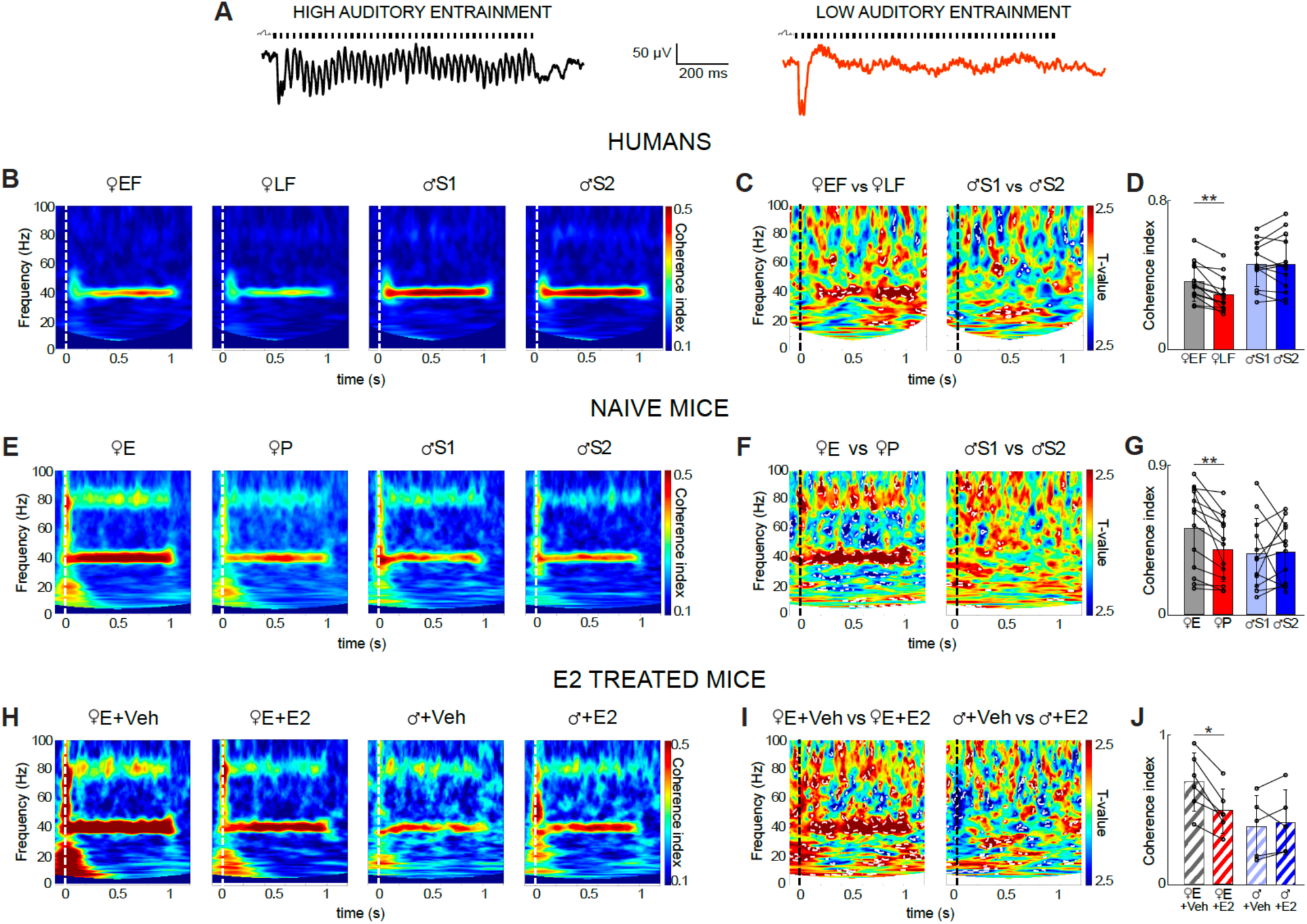
High estradiol levels are associated with lower inter-trial coherence during steady state stimulation at 40Hz across in mice and humans. [A] Examples of representative LFP traces illustrating high (left) and low (right) auditory entrainment during 40-Hz auditory steady state st mulation. [B] Time–frequency maps of inter-trial coherence in humans. The onset of auditory stimulation is indicated by a white dotted line. [C] Paired statistical maps showing differences in ITC between conditions in humans. Time– frequency regions exhibiting significant differences (P < 0.05) are outlined by white dotted lines; stimulus onset is indicated by a black dotted line. [D] Coherence index values corresponding to the stimulating frequency (40 Hz) in humans. Wilcoxon signed-rank: women EF vs LF: test z=3.1798, p = 0.0015; men S1 vs S2: z=-0.1048, p = 0.9165. [E] Time–frequency maps of inter-trial coherence in naïve mice. The onset of auditory stimulation is indicated by a white dotted line. [F] Paired statistical maps showing differences in ITC between conditions in naïve mice. Time– frequency regions exhibiting significant differences (P < 0.05) are outlined by white dotted lines; stimulus onset is indicated by a black dotted line. [G] Coherence index values corresponding to the stimulating frequency (40 Hz) in naïve mice. Wilcoxon signed-rank: females in estrus vs proestrus: z = 3.1798, p = 0. 0015; males S1 vs S2: z = −0.3922, p = 0. 6949). Only ASSR occurring with periods of immobility were analyzed. [H] Time–frequency maps of inter-trial coherence for each experimental group in E2 treated mice. The onset of auditory stimulation is indicated by a white dotted line. [I] Paired statistical maps showing differences in ITC between conditions in E2 treated mice. Time–frequency regions exhibiting significant differences (P < 0.05) are outlined by white dotted lines; stimulus onset is indicated by a black dotted line. [J] Coherence index values corresponding to the stimulating frequency (40 Hz) in E2 treated mice. Wilcoxon signed-rank: females in estrus + Veh vs + E2: z = 2.2014, p = 0.0277; males + Veh vs + E2: z = −0.6732, p = 0.5002. Only ASSR occurring with periods of immobility were analyzed. *P < 0.05, **P < 0.01. EF: early follicular; LF: late follicular; E: estrous; P: proestrus. Bar plots show mean ± SEM. Dots represent individual data. Number of human participants: women = 13; men = 13. Number of naive mice: females = 13, male = 12. Number of E2 treated mice: females = 6 male = 5.

Together, these findings indicate that E2 is sufficient to rapidly alter the temporal organization of sensory-evoked cortical activity, selectively reducing the consistency with which neural responses align to 40-Hz rhythmic auditory input.

### Estradiol reorganizes the temporal structure of neuronal firing during 40-Hz auditory entrainment

We next examined how the estrous cycle modulates stimulus-evoked spiking activity across different neuronal populations in female mice. We observed a pronounced estrous cycle-dependent modulation selectively affecting narrow-spiking (NS) neurons following a 40Hz, but not a 10Hz, auditory stimulation (Figure 4A-F, Supplemental Figure 9). Spike-density functions derived from peristimulus time histograms showed that NS neurons exhibited comparable initial responses at the 40Hz-stimulus onset across cycle stages but diverged markedly in their sustained activity (Figure 4B). During estrus, NS neurons rapidly decreased their firing rate following the initial response (0-0.1 s following stimulus onset), whereas during proestrus they show sustained firing throughout the 1-s stimulation period (Fig. 4F). Thus, the high-E2 state was characterized by enhanced sustained recruitment of putative fast-spiking interneurons during prolonged rhythmic stimulation.

**Figure 4.**
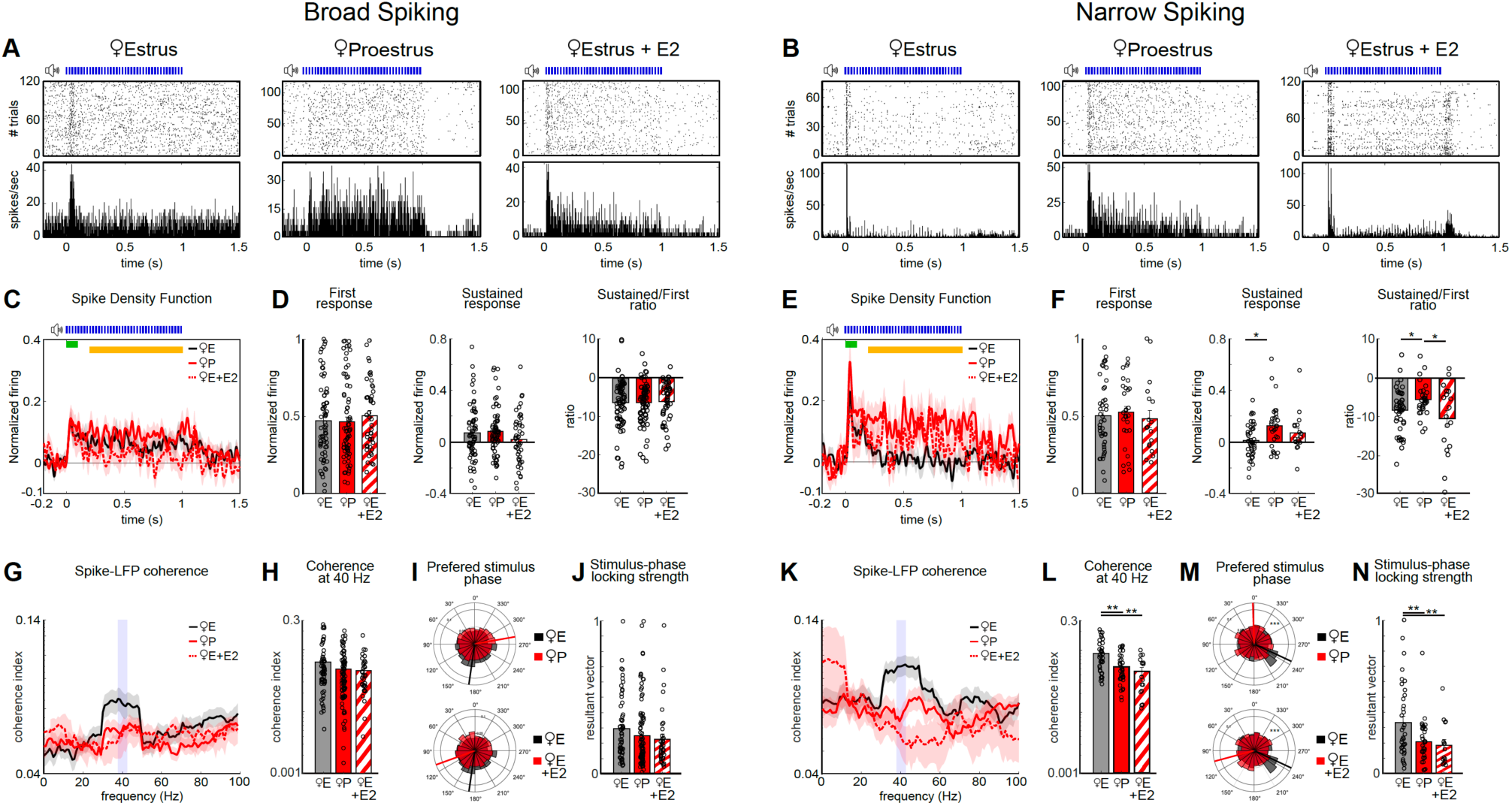
Estrous cycle modulates firing rate and spike–LFP coupling during 40 Hz auditory steady-state stimulation in mice. [A] Example raster plots (top) and peri-stimulus time histograms (bottom) for broad-spiking neurons during estrus (left), proestrus (middle) and estrus+E2 (right). [B] Example raster plots (top) and peri-stimulus time histograms (bottom) for narrow-spiking neurons during estrus (left), proestrus (middle) and estrus+E2 (right). [C] Spike density functions (shaded) for responsive broad-spiking neurons in female mice during estrus (black), proestrus (red) and estrus + E2 (dotted red). The early response window (0–100 ms post-stimulus) is indicated in green, and the sustained response window (200–1000 ms) in yellow. [D] Firing rates of broad-spiking neurons during the early and sustained response windows, and their ratio (sustained/early). Normalized firing rate shows no difference during early response (LMM, estrus vs proestrus: p = 0.6913; estrus vs estrus+E2: p = 0.6587), sustained response (LMM, estrus vs proestrus: p = 0.7818; estrus vs estrus+E2: p = 0.1547), ratio (LMM, estrus vs proestrus: p = 0.9854; estrus vs estrus+E2: p = 0.7583). [E] Spike density functions (shaded) for narrow-spiking neurons in female mice during estrus (black), proestrus (red) and estrus + E2 (dotted red). The early response window (0–100 ms post-stimulus) is indicated in green, and the sustained response window (200–1000 ms) in yellow. [F] Firing rates of narrow-spiking neurons during the early and sustained response windows, and their ratio (sustained/early). Normalized firing rate did not show differences between groups at the early response (LMM, estrus vs proestrus: p = 0.3903; estrus vs estrus+E2: p = 0.2671), but their sustained response (LMM, p = 0.0138) and ratio (LMM, p = 0.0332) were significantly different between estrus and proestrus. Acute estradiol during estrus did not affect sustained response (LMM, p = 0.1437) or the sustained/early ratio (LMM, p = 0.2222). [G] Spike–LFP coherence spectra for broad-spiking responsive neurons. [H] Spike–LFP coherence at 40 Hz of broad-spiking neurons shows no difference between groups (LMM, estrus vs proestrus: p = 0.1298; estrus vs estrus+E2: p = 0.2722). [I] Circular histograms show the distribution of spike phases relative to the 40-Hz stimulus cycle and the preferred firing phase between groups for broad-spiking neurons (Watson-U2 test, estrus vs proestrus: U=0. 1543, p=0.0950; estrus vs estrus+E2: U=0.1471, p=0.1097). [J] Strength of stimulus-phase locking, quantified by the resultant vector length in broad-spiking neurons (LMM, estrus vs proestrus: p = 0.3723; estrus vs estrus+E2: p = 0.1397). [K] Spike–LFP coherence spectra for narrow-spiking responsive neurons. [L] Spike–LFP coherence at 40 Hz of narrow-spiking neurons. LMM, estrus vs proestrus, p = 0.0016; estrus vs estrus+E2, p = 0.0090. [M] Circular histograms show the distribution of spike phases relative to the 40-Hz stimulus cycle and the preferred firing phase between groups for narrow-spiking neurons (Watson-U2 test, estrus vs proestrus: U=0.9139, p=2.9281*10^−8^; estrus vs estrus+E2: U=0.4284, p=4.2477 *10^−4^). [N] Strength of stimulus-phase locking, quantified by the resultant vector length in narrow-spiking neurons (LMM, estrus vs proestrus: p = 0.0059; estrus vs estrus+E2: p = 0.0020). *P<0.05, **P<0.01. E: estrous; P: proestrus. Bar plots show mean ± SEM. Dots represent individual neuron data. Broad Spiking cell number: estrus = 70, proestrus= 90, estrus + E2 = 36. Narrow Spiking cell number: estrus= 37, proestrus= 34, estrus+E2= 16. N female mice=13.

We next asked whether this increased firing was accompanied by changes in the temporal organization of neuronal activity relative to the ongoing 40-Hz oscillation. We quantified spike– LFP coherence during the entrained period of the stimulation (0.2–1 s). Consistent with our observations at rest, NS neurons exhibited significantly higher spike–LFP coherence at 40 Hz during estrus than during proestrus (Figure 4K, L), whereas no cycle-dependent difference was observed in BS neurons (Figure G, H). BS neurons showed a similar tendency toward reduced 40-Hz spike–LFP coherence during proestrus (Figure 4G, H); however, this effect was not significant when using the linear mixed-effects model, which accounts for variability across both cells and animals. Thus, despite their greater sustained firing during proestrus, NS neurons were less strongly coupled to the ongoing 40-Hz population rhythm. This reduced spike–LFP coherence could arise from reduced temporal precision of the underlying LFP oscillation, consistent with the lower 40-Hz ITC described above, from a reorganization of the temporal structure of neuronal spiking, or from a combination of both. To further examine the temporal organization of spiking independently of response magnitude, we assessed within-cycle phase locking using circular statistics. Watson’s U² test revealed significant differences in spike-phase distributions between estrous phases in the NS, but not in the BS, neuronal population (Fig. 4 I, M), indicating that their organization of spike timing within the oscillatory cycle differed according to hormonal state. Since these population-level circular analyses were performed on pooled spike-phase distributions, they did not weight individual neurons equally: neurons with higher firing rates contributed a larger number of spikes and could disproportionately influence the resulting population phase distribution. To control for this potential bias and assess phase locking at the level of individual neurons, we quantified each neuron’s phase preference using resultant vector length and compared these values between groups using linear mixed-effects models (Figure 4 J, N). This analysis revealed a significant cycle-dependent difference selectively in NS neurons, with stronger phase locking to the stimulus during estrus (figure 4N). Therefore, despite their enhanced sustained recruitment during proestrus, NS neurons showed reduced temporal coordination with both the 40-Hz stimulus and the concomitant LFP oscillations, indicating that NS neurons were particularly sensitive to estrous cycle-dependent modulation of both response magnitude and temporal organization.

Next, we examined stimulus-evoked spiking following acute E2 administration in females at the estrus phase. In contrast to the endogenous high-E2 state, acute E2 treatment did not increase the sustained firing rate of NS neurons, which remained comparable to that observed during estrus (Figure 4 E, F). However, E2 significantly reduced NS spike–LFP coherence at 40 Hz and decreased stimulus-phase locking strength, as shown by the resultant vector length (Figure 4 K-N). When analysing single neuron firing in response to a 10 Hz stimulation, we observed that both estrous cycle and acute E2 administration significantly affected spike-LFP coherence of NS neurons (Supplemental figure 9 K, L), while their stimulus-phase locking strength did not show significant changes (Supplemental figure N).

Together, these observations indicate that acute E2 administration is sufficient to rapidly reorganize the temporal coordination of putative fast spiking inhibitory neuron activity following high frequency stimulations, while other mechanisms, possibly engaged by prolonged E2 exposure or the influence of additional ovarian hormones, promote their enhanced sustained recruitment during proestrus.

## DISCUSSION

Despite 50 years of research on how circulating sex hormones influence behavior and cognitive function^5^, their effects on neuronal activity and network dynamics remain unclear. Furthermore, to what extent the ovarian cycle’s regulation of network states is evolutionarily conserved across species remains an open question. Here, we show that the ovarian cycle modulates aperiodic, but not oscillatory, network activity at rest and the temporal encoding of incoming sensory information at 40Hz in both mice and women. We further show that exogenous E2 administration rapidly increases the steepness of the aperiodic spectral slope, while simultaneously reducing the temporal precision of the auditory responses to fast rhythmic stimulations in female but not male mice.

One of our main findings is that E2-dependent reduction in conventional gamma-band power did not reflect a selective suppression of gamma oscillations, but rather a change in the broadband aperiodic component of cortical activity. Electrophysiological signals exhibit both periodic and aperiodic properties, and failure to dissociate these components can lead changes in the aperiodic background to be misinterpreted as changes in oscillatory power within predefined frequency bands^34^. Analyzing these two components together rather than separately may account for the conflicting findings in previous studies examining how the estrous cycle affects cortical oscillations^24,25^. Further, both components warrant investigation: while periodic oscillations are extensively associated with diverse physiological, cognitive, behavioral, and disease states^19,35–41^, growing evidence indicates that the aperiodic component is not neural noise, but rather encodes meaningful physiological information and dynamically changes with age, task demands and cognitive states^34,42–46^. In particular, the slope of the aperiodic power spectrum has been proposed as a proxy of changes in excitation/inhibition (E:I) balance ^28,30,47^, with steeper slopes reflecting a reduced E:I ratio. Here, we found that higher-E2 states were consistently associated with a steeper aperiodic spectral slope across species. Further, we demonstrated that exogenous E2 administration rapidly increased the steepness of the aperiodic spectral slope in female mice, thus suggesting that high E2 levels promote a shift toward greater inhibitory relative to excitatory synaptic activity^28,29,48^.

Supporting this hypothesis, narrow-spiking neurons (which are thought to be fast-spiking inhibitory interneurons) displayed higher spontaneous firing rates during proestrus, potentially indicating greater recruitment of local inhibition during high-E2 states. Fast-spiking interneurons exert powerful control over population activity, and changes in their recruitment or function can profoundly alter state-dependent and oscillatory network dynamics^49–51^. While we have not confirmed the specific cellular identity of narrow-spiking neurons, our data is consistent with a previous report showing that E2 increases the firing of fast-spiking GABAergic interneurons in rodent somatosensory cortex^10^ and that local estrogen synthesis regulates fast spiking interneuron activity, inhibitory transmission, and network oscillations in the female hippocampus^52^.

A second main finding is that a high E2 estrous state was associated with higher firing frequency, but reduced spike–LFP coherence of putative fast spiking neurons, both at rest and during a fast rhythmic auditory stimulation, indicating that this increased activity was less precisely organized relative to the ongoing population-level field activity. In parallel, entrainment at 40Hz was attenuated, suggesting that the temporal organisation of narrow-spike neurons may contribute to the temporal precision of auditory responses during this fast rhythmic stimulation. This dual role of E2 may not be exclusive to the mammalian cortex. In fact, in the songbird auditory forebrain locally synthesized E2 reduces intrinsic excitability, while increasing the temporal reliability of neuronal firing^13^. However, these effects are heterogeneous across neuronal populations, indicating that E2 does not simply shift auditory circuits toward excitation or inhibition but instead reconfigures the cellular interactions that determine population dynamics^13^. By dissociating neuronal firing rate from temporal coordination, E2 could reshape how individual neurons participate in population-level representations of auditory inputs. Of note, exogenous E2 administration rapidly reduced spike-LFP coherence, without changing the firing frequency of NS neurons. E2 can both rapidly induce ion fluxes and the activation of protein kinases across the plasma membrane, independently of protein synthesis (non-genomic effects), and change gene transcription (genomic effects) on a longer time scale. Because electrophysiological recordings occurred shortly after E2 administration (∼15 min), the altered network dynamics and spike–field coordination are likely driven by non-genomic mechanisms. Conversely, shifts in baseline interneuron excitability may depend on longer-term, sustained elevations of E2 during proestrus. It is also possible that the increased interneuron firing rate observed during proestrus may depend on additional hormonal or circuit adaptations.

The third main finding is that the ovarian cycle modulates the temporal encoding of ongoing sensory input at 40Hz in both mice and women, with reduced entrainment during high E2 phases across species. Because auditory entrainment has been proposed to temporally compartmentalize incoming sensory inputs^32,33^, we hypothesize that E2-dependent changes in temporal synchronization may shift the way auditory cortical networks sample acoustic information. During low-E2 states, greater temporal precision of fast-spiking activity may drive LFP stronger intertrial coherence and reinforce the dominant oscillatory structure, favoring the reliable encoding of inputs aligned with its optimal phase and thereby supporting the representation of global temporal regularities (beat perception). Conversely, during high-E2 states, increased spontaneous firing but reduced temporal coupling of fast-spiking neurons may distribute inhibition more broadly in time and increase LFP trial-to-trial phase variability. This temporal desynchronization could broaden the windows over which incoming inputs are integrated, allowing information arriving outside the preferred phase of the dominant rhythm to be represented and potentially increasing sensitivity to subtle variations in the acoustic environment (pitch perception). This interpretation is consistent with behavioral evidence suggesting cycle-dependent differences in auditory perception. Auditory beat perception has been reported to vary across the menstrual cycle, with better binaural and physical beat perception during menstruation, suggesting that low-hormone states may favor the representation of periodic auditory structure^53^. Conversely, auditory discrimination, including frequency-and pitch-related measures, has been reported to improve around the high-E2 periovulatory phase^54,55^, potentially reflecting greater sensitivity to fine acoustic details. Together, these observations raise the possibility that E2-driven changes in the temporal organization of fast-spiking activity shift cortical processing between distinct sampling strategies: a temporally constrained mode that favors precise representation of rhythmic regularities and a more flexible mode that broadens the integration of incoming sensory information. Future studies directly linking hormonal fluctuations, cortical temporal dynamics, and auditory behavior will be required to determine whether this proposed mechanism contributes to cycle-dependent changes in sensory perception.

In summary, this work identified evolutionary conserved effects of the ovarian cycle on network states and auditory responses evoked by fast rhythmic stimulation. It further highlights narrow-spiking neurons as potential target of E2 action. The molecular and synaptic mechanisms underlying the differential effects of E2 on these neurons firing rate vs LFP-spike coherence remain to be investigated. Furthermore, we cannot exclude that other neuron types, for example somatostatin-expressing interneurons, likely included in the broad spiking neuronal population, are modulated by E2 levels. Finally, how the estrous’ cycle associated changes in aperiodic activity modulate state related neural dynamics needs to be investigated. Nevertheless, since EEG-based signatures are extensively used as biomarkers in fundamental studies and clinical trials for neurodevelopmental and psychiatric disorders, our findings indicate that the ovarian cycle and E2 represent biologically meaningful sources of neuromodulation that should be considered in experimental design and interpretation. More broadly, many brain disorders show marked sex differences in prevalence, clinical expression, and disease trajectory^56–59^. By identifying conserved, hormone-sensitive electrophysiological signatures across mice and humans, our study provides a cross-species framework in which the contribution of ovarian hormones to these differences can be experimentally investigated.

## MATERIALS AND METHODS

### MICE

**Animals.** Experimental procedures involving mice were approved by the Comité Institutionnel de Bonnes Pratiques Animales en Recherche (CIBPAR) of the Research Center of Sainte-Justine Hospital in accordance with the principles published by the Canadian Council on Animal Care. A total of 25 mice were used in this study, including 13 females and 12 males on a C57BL/6 background. Mice were implanted at postnatal day (p)70–90 and recorded between p80 and p120. All animals were housed with two to five mice/cage from weaning until surgery, after which they were housed singularly for the duration of the experiment. Animals were kept on a 12 h/12 h light/dark cycle and provided with nesting material and food and water ad libitum. All experiments were performed during the light period under constant mild luminosity (60 lx). None of the recorded animals were excluded from the final analysis.

**Estrous cycle phase classification.** Vaginal smear cellular composition was used to determine estrous cycle phases based on the relative proportions of leukocytes, nucleated epithelial cells, and cornified cells. Vaginal smear samples were taken after recording. Because the estrous cycle is a continuous process, transitions between phases do not present sharp boundaries but rather reflect gradual shifts in cellular composition; therefore, phase classification should be considered an approximation of these dynamic changes. This study includes data from proestrus (Supplemental figure 2, red square), encompassing: early proestrus, defined by the emergence of nucleated epithelial cells (>∼10% of total cells, with the remainder primarily leukocytes), and late proestrus, characterized by a predominance of nucleated cells (∼85%) with a small proportion of cornified cells (∼10%) and occasional leukocytes (∼5%). It also includes estrus (Supplemental figure 2, black square), comprising: early estrus, when cornified cells exceed ∼15% of total cells (with the remaining cells being nucleated), and late estrus, marked by the near-complete dominance of cornified cells with the initial reappearance of leukocytes.

**Estradiol and vehicle injection.** After being recorded during multiple estrous cycles, female mice at the end of the estrus phase, received an intraperitoneal (i.p.) injection of either vehicle (saline) or water-soluble 17β-estradiol (Sigma-Aldrich, E4389; 45 mg/g). Mice received both vehicle and 17β-estradiol injection in randomized order at least one week apart. 17β-estradiol was dissolved in saline and administered at a dose of 20 µg/kg body weight. Injections were performed 15 min before the start of the resting-state recording and 25 min before the auditory steady-state response paradigm.

**Microdrive preparation**. Each tetrode was fabricated out of four 10-μm-diameter tungsten wires (H-Formvar insulation with Butyral bond coat California Fine Wire Company, Grover Beach, CA) that were twisted and then heated to bind them into a single bundle. Each tetrode was then assembled to an individually moving shuttle and they were then assembled within a Microdrive structure in which a screw individually move each shuttle-tetrode. The tips of the tetrodes were freshly cut the day of implantation and gold-plated to reduce the impedance to < 400 kΩ. The use of tetrodes provides not only LFP recordings but also high-density spiking activity, allowing for the characterization of single-unit discharges. The 3D-printed microdrives used were manufactured by Axona (Axona Ltd.; http://www.axona.com/).

**Surgery.** Mice were implanted at postnatal day (p)70–90 and recorded between p80 and p120. Chronic implants were performed under deep anaesthesia, induced by an intraperitoneal injection of Ketamine/Xylazine/Acepromazine cocktail (80 mg/kg, 15 mg/kg, and 2 mg/kg respectively). Body temperature was maintained at 37°C using a feedback-controlled heating pad. Analgesics were given locally subcutaneously (lidocaine 2%, Aspen Pharmacare, Canada) and an intraperitoneally dose of 30 mg/kg (buprenorphine, 0.3 mg/mL, Vetergesic Multidose, Ceva) 5 min before surgery. Mice were head-fixed in a stereotaxic frame (Kopf Instruments, USA). During the entire surgery, eyes were protected and hydrated using Optixcare Eye Lube (CLC Medica, Canada). The skull was exposed, cleaned, and treated with an activator (Metabond Quick, Parkell, USA) to increase bone porosity. The coordinates of the targeted area were marked with a permanent marker before fixing the skin around the exposed skull with Vetbond tissue adhesive (3M, USA). Then a first layer of dental cement (Metabond Quick, Parkell, USA) was applied to cover the whole exposed skull except the marked coordinates. Craniotomies were then performed on the right side of the primary auditory cortex (A1, −2.5 mm posterior to Bregma and 4 mm lateral to the midline) and on the contralateral frontal lobe (area chosen to place the ground-and-reference wire). Four independently movable tetrodes were then placed vertically at 0.5 mm of depth, immediately above A1, layer V. A drop of silicone elastomer (WPI, USA) was applied around the tetrodes to let tetrodes free for movement. The microdrive apparatus and the ground-and-reference wire were then daubed with dental acrylic (Ortho-Jet Powder and Jet liquid, Lang Dental, USA) to encase the microdrive and anchor it to the skull.

**Electrode’s placement verification.** Electrode placement was confirmed at the end of each experiment via post-hoc immunohistochemistry (Supplemental Figure 2). Following the conclusion of data acquisition mice were perfused transcardially with saline followed by 4% paraformaldehyde in phosphate buffer (PB, 0.1 M, pH 7.2). Brains were post-fixed with 4% paraformaldehyde overnight and the implanted recording electrodes were carefully removed. Subsequently the brains transferred to a 30% sucrose solution in PBS for 48 h. Coronal sections of 40 µm thickness were obtained at optimal cutting temperature using a cryostat (Leica Microsystems VT100). Brain sections were blocked in 10% normal goat serum and 1% Triton X-100 for 2 h at room temperature. Slices were then incubated for 48 h at 4°C with guinea pig anti-GFAP (1:3000, Synaptic Systems, Cat. No. 173004). This step was followed by incubation with secondary antibodies for 2 h at room temperature to visualize primary antibodies. The secondary antibodies used were Alexa-Fluor conjugated 647 (1:1000, Life Technologies). After rinsing in PBS (three times), brain sections were stained with DAPI (1:1000 in PBS) for 15 min at room temperature to visualize cell nuclei. Brain sections were then rinsed another three times in PBS and mounted in Vectashield mounting medium (Vector). Whole-brain slices were first imaged using the AxioScan7 Digital Slide Scanner (Zeiss) at 10x objective to quickly localize the sections where the electrode was positioned. Subsequently the selected slices were imaged on a wide-field fluorescent microscope (Leica DMi8) using 10× and 20× objectives (Supplemental Figure 1). For each slice, 10× images of the entire slice were acquired as a series of overlapping tiles and stitched automatically by the microscope software to generate a single high-resolution image. Single-plane images were captured for each slice. Higher-magnification 20× images were obtained at the auditory coordinates to confirm electrode placement.

**Data Acquisition.** LFP recordings were performed using an open-ephys GUI platform (https://open-ephys.org/). Data were acquired at 20 KHz with a custom-made headstage using Intan Technologies RHD Electrophysiology Amplifier chip (https://intantech.com/products_RHD2000.html). Custom-designed auditory and visual trigger generator devices sent triggers directly to the recording system, assuring the precise timing of stimuli presentation. Starting 1 week after surgery, animals were habituated daily for 4 days to an open field (15–20 min each time), before starting recordings for auditory stimulation protocols from Day 5. We used a dimly illuminated open field environment (45 × 35 cm), surrounded by 60-cm-high transparent walls, equipped with video monitoring. The walls were sprayed and wiped clean with odor-free Peroxigard (Virox Technologies) 15 min before the introduction of each animal. Two speakers were placed ∼10 cm from the walls of the open field. Resting state activity was recorded at the beginning of each session for a period of 10 min, while mice were freely exploring the open field. Data acquisition during auditory stimulations was conducted for 10– 15 min, a total of three to six times, on alternate days.

**Auditory Steady State Protocol.** Auditory stimuli were presented at an intensity of 70–75 dB SPL. A total of 240 click trains were delivered, comprising 120 trials at 10 Hz and 120 trials at 40 Hz, presented in an alternating sequence. Each click train lasted 1 s, with an inter-trial interval randomly jittered between 2 and 5 s. Individual clicks consisted of 5 kHz tones with a duration of 5 ms. The entire stimulation protocol lasted approximately 13 min.

**Data Processing.** LFP signals were down-sampled to 2000 Hz, and a bandpass filter (0.5–150 Hz) was applied. A notch filter (59.5–60.5 Hz) was also applied to remove residual 60 Hz noise contamination. For resting state analysis data was segmented in immobility periods, that were defined by establishing a velocity threshold of the animal body centre at < 2 cm/s for at least 2 seconds. To analyze the evoked responses, data was segmented into 2500 ms (500 ms pre-stimulus and 2000 ms post-stimulus onset) periods. Trials with an intra-channel average higher than four times the total trial standard deviation (SD) were tagged, visually inspected and removed. Only trials in which the onset of the stimuli (−100 +200) overlapped with an immobility period were taken into consideration. Only sessions containing more than 20 clean trials during immobility were kept for further analyses.

### HUMANS

**Participants.** Experimental procedures involving human participants were approved by the ethics, scientific, and administrative committees of CHU Sainte-Justine. Written informed consent was obtained from all participants prior to inclusion in the study. Participants aged 20–40 years of both sexes were recruited. Data were collected from 15 regularly cycling women and 13 age-matched neurotypical male control subjects with comparable levels of education. The mean age was 31.38 years for women and 32.54 years for men. Only neurotypical individuals without a personal or family history of neurodevelopmental disorders (e.g., intellectual disability or autism spectrum disorder) were included. None of the participants were taking medication or receiving hormonal therapy at the time of the study.

Women were included if they reported regular menstrual cycles lasting 28–30 days and were not using hormonal contraceptives. Two female participants were excluded: one due to a technical issue during data acquisition (less than 30 trials remined after artifact rejection) and another due to thyroid-related medication not reported during the first visit. Therefore, a total of 13 women were included in the final analyses. Women were recorded twice within their cycle: during the first 48 h (early follicular phase), when hormone levels are lowest, and approximately 10 days later (late follicular phase), when estradiol levels typically peak ^60^. All comparisons were performed within subjects using paired analyses.

**EEG Recordings and Sensory Stimulation Protocols**. EEG recordings were conducted in an electrically shielded and sound-attenuated room at CHU Sainte-Justine. Signals were acquired using a 128-channel Geodesic Sensor Net (Electrical Geodesics, Inc.) connected to an EGI Net Amps 300 amplifier and recorded with NetStation software (version 4.5.4). Data were sampled at 1000 Hz using the vertex electrode as an online reference and filtered with a lower bound of 0.5 Hz and a high cut-off at 100 Hz during data acquisition. Electrode impedances were maintained below 40 kΩ. For the analyses reported here, we focused on Cz and the surrounding central electrodes, as auditory cortical ERPs, particularly the N1–P2 complex, are typically prominent over frontocentral scalp regions and show maximal or near-maximal amplitudes at the vertex (Cz). The COM electrode served as the analogue reference during EEG acquisition.

Resting-state recordings lasted 4 minutes and consisted of an eyes-open cross-fixation task, during which participants were instructed to look at a cross fixed in the center of the screen located approximately 60 cm from them. Lights were kept on during resting-state recordings and turned off during auditory stimulation. Auditory stimuli were delivered via two lateral speakers (BX5a, M-Audio) positioned approximately 30 cm from the participants’ ears. Stimuli were generated using E-Prime 2.0 (Psychology Software Tools, Inc.) on a Dell Optiplex 790 computer. During auditory stimulation, participants were instructed to passively listen to the sounds while watching a muted movie without subtitles. The auditory steady-state stimulation protocol was identical to that used in mice (see above).

**Data Processing.** Original data saved in mff format was converted to set using eeglab. Data processing was performed using HAPEE open source matlab codes^61^. Channel rejection was performed automatically using HAPPEÈs default configuration. A bandpass filter (0.5–150 Hz) and a notch filter was applied. Blink artefacts and remaining muscular artefacts were removed using automatic independent component analysis (ICA). Data were re-referenced to an average reference (E1001). For the analysis of event-related potential, data were segmented from −500 to 1500 ms relative to the stimulus onset, and the signal was baseline-corrected from −200 to 0 ms. Segment rejection was based on both similarity and amplitude (−150 +150 mV). Auditory processing in humans is reported to peak in vertex and midline electrodes^62,63^, thus we selected the midline central (Cz) electrode and the five electrodes surrounding it, as the region of interest for our analyses on auditory perception. Data from these 6 electrodes were then averaged. Only one participant was excluded (as mentioned before) because one session contained less than 30 trials.

### EEG/LFP DATA ANALYSIS

Data analysis and quantification were performed using custom MATLAB (The MathWorks Inc.) codes, available upon request. Animal behavior was analyzed using DeepLabCut^64^. Body center was extracted and used to measure mice locomotor activity.

**Power Spectral Density in Mice and Humans.** Resting state spectral activity was analyzed during resting periods in the absence of auditory stimulation using two complementary approaches. First, conventional power spectral density (PSD) was estimated using the Welch method implemented in MATLAB to quantify power within predefined frequency bands, adapting code from https://github.com/elifesciences-publications/Miri-Vinck-et-al. The estimated power spectrum was normalized to the total signal power and plotted on a logarithmic scale for visualization, facilitating comparison with previous studies in the field^65^.

Second, to determine whether differences observed in conventional frequency-band power reflected changes in periodic oscillatory activity or in the underlying broadband spectral background, power spectra were parameterized using the Fitting Oscillations & One-Over-F (FOOOF) approach^34^, as implemented in FieldTrip. For this analysis, spectra were estimated using a multitaper fast Fourier transform (mtmfft; 1-Hz spectral smoothing, 5-s zero-padding) over the 0–100 Hz frequency range. The original power spectrum was decomposed into its periodic and aperiodic components. The fitted aperiodic component was subtracted from the original power spectrum to obtain the oscillatory spectrum, allowing frequency-specific oscillatory power to be quantified independently of broadband spectral changes. The aperiodic spectral slope was quantified from the FOOOF-derived aperiodic component by fitting a linear regression to the spectrum in log–log coordinates. The slope and intercept were extracted for each recording and averaged across sessions for each subject or animal, with more negative slope values corresponding to a steeper aperiodic spectrum.

**Time Frequency Spectrograms in Mice and Humans.** Time-frequency spectrogram analysis allows exploration of the power of different frequency bands over time. We used the complex Morlet’s wavelet transformation to provide power maps in time and frequency domains^66^. The simplified mathematical expression for applying this specific wavelet convolution is:

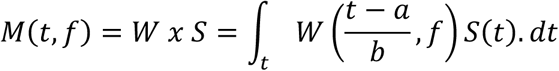

where M(t,f) is a matrix of complex values (vectors) for a given time (t) and frequency (f). S is the signal as a function of time (t) and W corresponds to Morlet’s wavelet, which is a complex exponential (Fourier) with a Gaussian envelope that undergoes a series of translations (a) and dilations (b) dependent on the frequency (f). Wavelet cycles increase linearly with frequency, beginning at 2 Hz [2,0.5]. Baseline correction was applied using the 500 ms prior to the stimuli onset. This analysis was performed using the EEGLAB toolbox (v.13.6.5b) (La Jolla, CA, USA) ^67^, in particular using the function newtimef.

**Inter-trial Coherence.** Analogous to the phase-locking value, inter-trial coherence allows assessment of the strength of phase coherence across trials in temporal and spectral domains^68^. The inter-trial coherence computation uses only the phase of the complex values given by Morlet’s wavelet transform. Inter-trial coherence measures phase coupling across trials at all latencies and frequencies and is defined by:

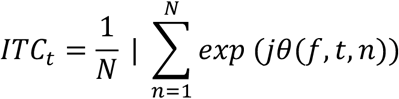

Where *jθ*(*f*, *t*, *n*) represents the phase for a given frequency (f), time point (t) and trial (n). The obtained values are always defined between 0 and 1. Phase-locking values close to 1 indicate strong inter trial phase-locking, thus representing evoked activity while scores closer to 0 indicate high inter trial phase variability^69,70^.

### SPIKING ACTIVITY ANALYSIS IN MICE

Offline processing of electrophysiological data began with the extraction of spikes from the high-pass filtered signal (800–5000 Hz) of recordings sampled at 20 kHz. Action potentials exceeding 5 standard deviations above the mean baseline were detected. Spike waveforms were then characterized by feature extraction, and putative neuronal units were identified using principal component analysis followed by automatic clustering using klustakwik^62^ (http://klustakwik.sourceforge.net/). Identified clusters were manually refined with a graphical cluster-cutting program^71^. Only units exhibiting a clear refractory period (< 2 ms) in their auto-correlograms and well-isolated cluster boundaries were included in subsequent analyses.

**Firing rate.** Peri-stimulus time histograms (PSTH) and spike density function (SDF) were performed using the mlib toolbox by Maik Stüttgen (https://de.mathworks.com/matlabcentral/fileexchange/37339-mlib-toolbox-for-analyzing-spike-data). PSTHs were calculated using a bin size of 4 ms. SDF were calculated by applying a Gaussian distribution over the PSTH using a kernel of 2 ms width. Firing rate was evaluated by classifying SDF profile into two components, early (0-100 ms following stimuli onset) and sustained (200-1000 ms following stimulus onset) response. We then identified three types of neurons, based on their firing behavior following the stimulus presentation: neurons that respond by increasing their firing (responding-excited), neurons that respond by decreasing their firing (responding-inhibited), and neurons whose firing does not change (non-responding neurons). This categorization was made by calculating the modulation index between their firing rate during baseline and upon stimulation by applying the formula:

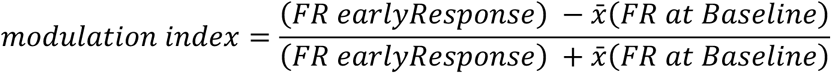

In this study, analyses were restricted to responsive neurons, defined by a modulation index > 0.2, as these constituted the majority of recorded units and provided sufficient statistical power. SDF shown here correspond to the average of all neurons’ SDF for each group. Sustained/Early ratio was calculated as:

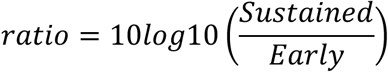

**Spike–LFP Coherence and Phase Preference**. Spike–LFP coherence was computed using the Chronux multitaper function coherencycpt^72^, which estimates the normalized cross-spectrum between the continuous LFP signal and spike times treated as a point process. Coherency was calculated as

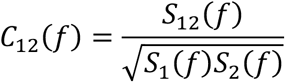

where S_12_(f) denotes the cross-spectrum between the LFP and spike train, and S_1_(f) and S_2_(f) are their respective auto-spectra. The coherency C_12_(f) is complex-valued, encoding both magnitude and phase relationships across frequencies. Coherence magnitude was defined as ∣C_12_(f)∣, and phase as:

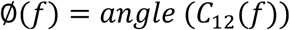

Spectral estimates were obtained using the multitaper method over a frequency range of 0–100 Hz, within a post-stimulus analysis window of 0.1–1 s, and using zero-padding. Tapers were defined as [TW,K]=[10,19], where TW is the time–bandwidth product controlling spectral concentration, and K is the number of orthogonal Slepian tapers used for estimation. This relatively high time– bandwidth product increases spectral smoothing and reduces variance, which is particularly important for stabilizing coherence estimates involving spike trains. Averaging was performed across tapers and trials.

In parallel, to assess the temporal organization of neuronal firing relative to the external 40-Hz auditory stimulus independently of the LFP, we quantified each spike’s phase within the 40-Hz stimulus cycle. Stimulus phase was calculated from spike timing relative to stimulus onset, with each 25-ms cycle corresponding to 0–2π radians. Spike phases were first pooled across neurons and trials to construct circular histograms and compare population-level preferred stimulus-phase distributions using circular statistics (Fig. 4I,M). In addition, stimulus-phase locking was quantified at the single-neuron level by calculating the resultant vector from the distribution of spike phases for each neuron. The angle of the resultant vector represented the neuron’s preferred stimulus phase, whereas its length provided a measure of stimulus-phase locking strength, ranging from 0 (no consistent phase preference) to 1 (all spikes occurring at the same phase of the stimulus cycle; Fig. 4J,N). Thus, whereas spike–LFP coherence quantified the relationship between neuronal firing and the ongoing LFP oscillation, these complementary circular analyses quantified the preferred phase and consistency of neuronal firing directly relative to the 40-Hz auditory stimulus.

### STATISTICAL ANALYSIS

Statistical analyses were selected according to the structure of each dataset. Paired comparisons of exploratory behavior and LFP/EEG measures were performed using two-tailed Wilcoxon signed-rank tests for both mouse and human datasets. This non-parametric paired approach was used to compare repeated measurements obtained from the same animals or participants across experimental conditions or hormonal states.

For single-unit spiking analyses, linear mixed-effects models (LMMs) were used to account for the hierarchical structure of the data, with multiple neurons recorded from the same animal. Experimental condition was included as a fixed effect and subject identity as a random effect, thereby accounting for the non-independence of neurons recorded within the same subject. This approach prevented animals contributing larger numbers of recorded neurons from disproportionately influencing group-level effects and ensured that statistical inference reflected variability across subjects rather than treating individual neurons as independent biological replicates.

Differences in circular spike-phase distributions were assessed using Watson’s U² two-sample test. This non-parametric circular test was used to determine whether the distributions of spike phases relative to the 40-Hz stimulus differed between experimental conditions. Because these population-level analyses were performed on pooled spike-phase distributions, complementary single-neuron analyses of stimulus-phase locking strength, quantified by resultant vector length, were analyzed using LMMs to account for the nesting of neurons within subjects.

Statistical tests were two-tailed, and differences were considered statistically significant at p < 0.05.

## ACKNOWLEDGEMENTS

We are grateful to all members of our lab for insightful data discussion. We would like to thank Dr Bénédicte Amilhon and Dr Guillaume Ducharme, Kristian Agbogba and Mikael Valton Charette for their technical assistance, the Comité Institutionnel de Bonne Pratiques Animales en Recherche (CIBPAR), all the personnel of the animal facility of the Research Center of CHU Sainte-Justine (Université de Montreal), Compute Canada, Elke Kuester-Shoeck and the CHUSJ Imaging platform (PIM) for their instrumental technical support. G.DC was supported by the Canadian Institutes of Health Research (CIHR, PG 19786), Natural Sciences and Engineering Research Council of Canada (NSERC, DG-2022-04444) and Fonds UdeM pour le partenariat CHU Sainte-Justine - Institut Imagine en épilepsie de l’enfant. M.I.C-M and A.C.C received scholarships from the Canadian Neurodevelopmental Research Training (CanNRT) Platform. M.I.C-M received scholarships from the Savoy Foundation. A.C.C and B.C. received scholarships from CIHR. A.C.C. received a scholarship from NSERC, funding reference #456461541. S.L was supported by Canada Research Chair Pediatric Neurodiversity (CRC-2024-00030), Fonds de recherche du Québec Programme AUDACE (337467), NSERC (DG-2023-04848), Fondation de la Recherche Pédiatrique.

