## Supplemental Figures for "The ovarian cycle and estradiol modulate cortical network state and auditory entrainment in mice and women"

Maria Isabel Carreño-Muñoz et al.

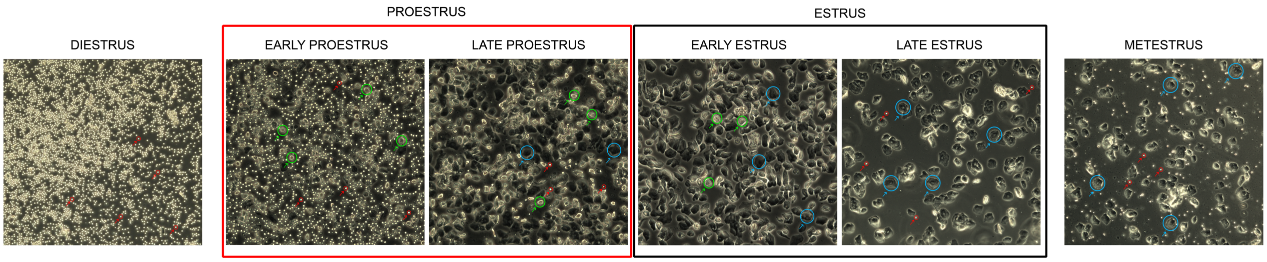


**Supplementary Figure 1. Vaginal smear cytology across estrous cycle stages in mice.** Estrous cycle stages were determined from unstained vaginal smear cellular composition, based on the relative proportions of leukocytes, nucleated epithelial cells, and cornified cells. As the estrous cycle is continuous, transitions between stages lack clear boundaries and instead reflect gradual shifts in cell proportions. Data include proestrus (red), spanning early (emergence of nucleated cells, >~10%) to late (predominantly nucleated cells ~85%, with ~10% cornified cells and occasional leukocytes), and estrus (black), spanning early (cornified cells >~15%) to late stages (complete dominance of cornified cells with initial reappearance of leukocytes). Representative images of diestrus and metestrus are also shown to illustrate the transition from the preceding phase (diestrus) and toward the subsequent phase (metestrus).

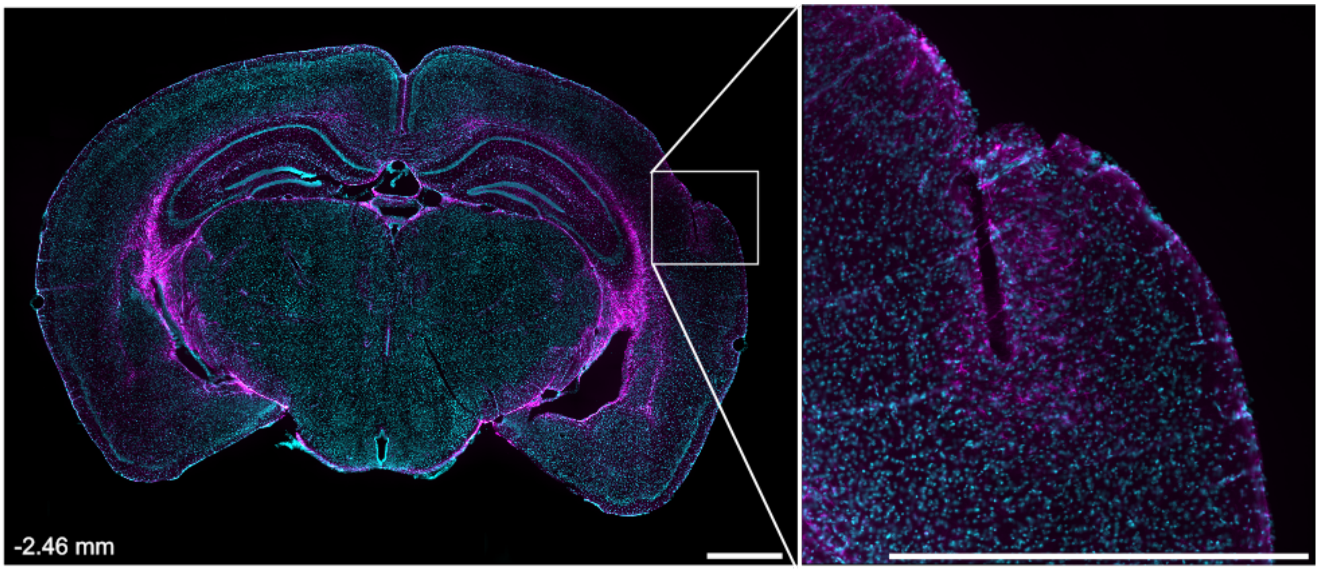


**Supplemental Figure 2. Electrode bundle positioning**.

Whole brain section (left) and high magnification (right) images showing electrode position for A1. Cell nuclei are labelled by DAPI (cyan) and astrocytes are labelled by GFAP (pink). Coordinates from bregma are indicated on the lower left corner. Scale bars: 1mm.

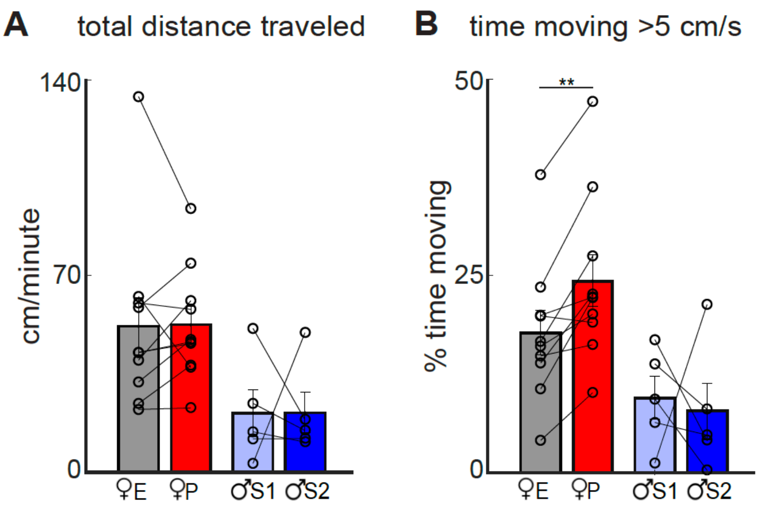


**Supplementary Figure 3. Locomotor activity by sex and estrous cycle phase.**

[**A**] Total distance traveled during 10 min of exploration in a familiar open field by female mice at estrus (E) and proestrus (P) (Wilcoxon signed rank test for paired samples z=0.6625, $p$ = 0.5076) and by male mice across two sessions (S1 and S2) (Wilcoxon signed rank test z=0.4045, $p$ = 0.6858).

[**B**] Total time spent in active displacement (movement speed >5 cm/s) during the same session by female mice during estrus and proestrus (Wilcoxon signed rank test z=-2.7011, $p$ = 0.0.0069) and by male mice (Wilcoxon signed rank test z=0.6742, $p$ = 0.5002). **$P$<0.01.

E: estrous; P: proestrus. Bar plots show mean. Dots represent individual mouse data. Number of mice: male = 5; females = 10.

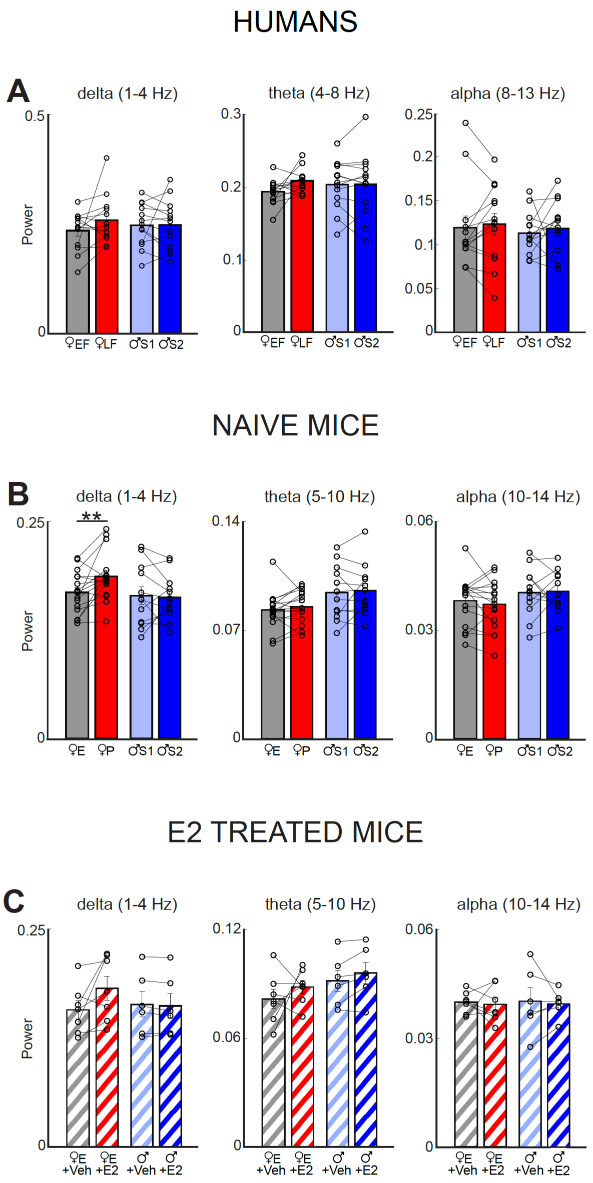


**Supplementary Figure 4. Baseline delta, theta, and alpha power across estradiol conditions in humans and mice.**

[**A**] Baseline power in the delta, theta, and alpha frequency bands in humans. Women were compared between the early follicular (EF; low E2) and late follicular (LF; high E2) phases of the menstrual cycle, while men were compared between two recording sessions separated by 10 days (S1 vs S2). Power was calculated from 4-min resting-state, open-eyes EEG recordings. Wilcoxon signed-rank test: delta band (women EF vs LF: $z = -1.5025$, $p = 0.1330$; men S1 vs S2: $z = 0.9435$, $p = 0.3454$), theta band (women EF vs LF: $z = -1.7122$, $p = 0.0869$; men S1 vs S2: $z = -0.1048$, $p = 0.9165$), alpha band (women EF vs LF: $z = -0.0524$, $p = 0.6002$; men S1 vs S2: $z = 0.9435$, $p = 0.3454$).

[**B**] Baseline power in the delta, theta, and alpha frequency bands in naïve mice. Females in estrus (low E2) were compared with females in proestrus (high E2), while males were compared between two recording sessions in an already familiarized open field (S1 vs S2). Power was calculated from immobility periods (>2 s) during the first 10 min of baseline recordings in freely behaving mice. Wilcoxon signed-rank test: the delta band (females in proestrus vs estrus: z=-2.8966 p=0.0038; males S1 vs S2: z=-0.5601 p=0.5754), theta band (females in proestrus vs estrus: z=-0.4544 p=0.6496; males S1 vs S2: z=0.8402 p=0.4008), alpha band (females in proestrus vs estrus: z=0.5071 p=0.1398; males S1 vs S2: z=-0.1400 p=0.8886).
[**C**]. Baseline power in mice following acute estradiol (E2) administration. To assess the acute effect of E2 on baseline oscillatory activity, female mice in estrus and male mice received either vehicle or E2, and delta, theta, and alpha power was compared between vehicle- and E2-treated conditions. Wilcoxon signed-rank test: delta band (females in estrus+Veh vs estrus+E2: z=-1.8593 p=0.0630; males+Veh vs males+E2: z=1.3628 p=0.1730), theta band (females in estrus+Veh vs estrus+E2: z=-0.6761 p=0.4990; males+Veh vs males+E2: z=-1.7821 p=0.0747); alpha band (females in estrus+Veh vs estrus+E2: z=1.4767 p=0.6121; males+Veh vs males+E2: z=-0.1048 p=0.9165).

Bar plots show mean ± SEM. Dots represent individual data. EF: early follicular; LF: late follicular; E: estrous; P: proestrus. Number of human participants: men = 13; women = 13. Number of naïve mice: females = 16, males =12; number of treated mice: females = 7, males =6.

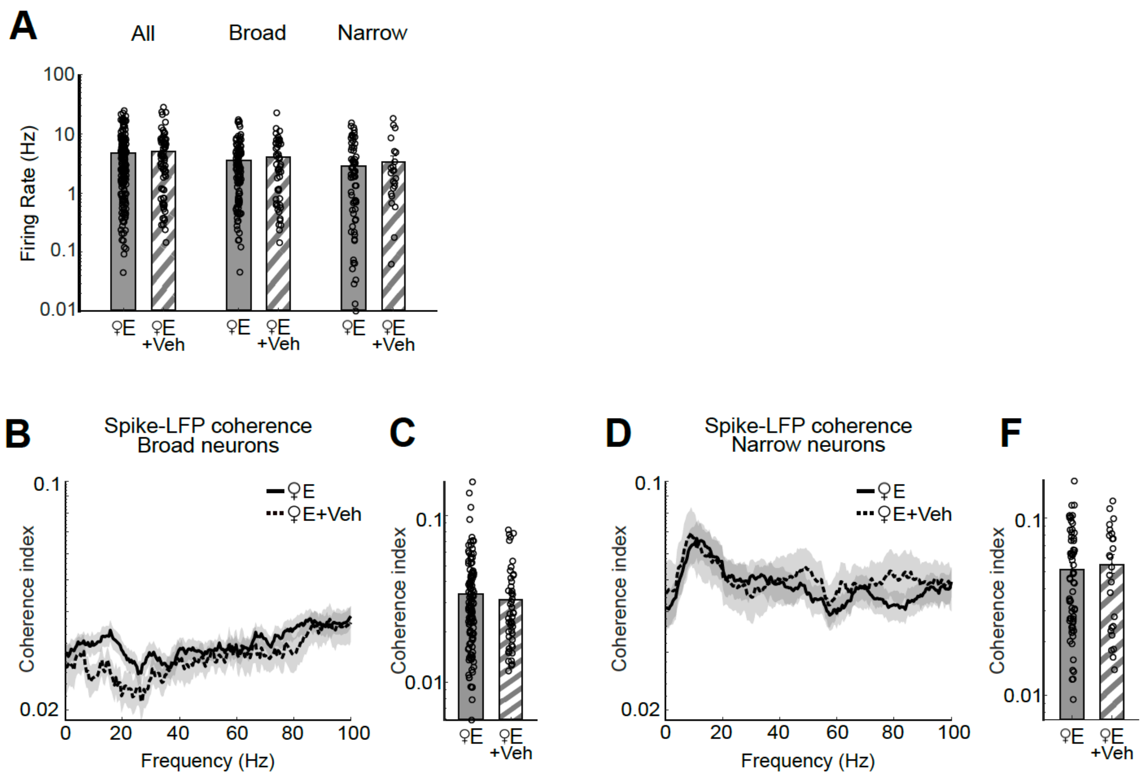


**Supplementary Figure 5. Vehicle injection does not affect spontaneous firing rate or spike-LFP coherence during immobility.**

[**A**] Mean firing rate during immobility periods (>2 s) within the first 10 min of exposure to a familiar open field. LMM, all neurons (females in estrus vs estrus+Veh: $p = 0.3888$), broad neurons (females in estrus vs estrus+Veh: $p = 0.2009$) and narrow neurons (females in estrus vs estrus+Veh: $p = 0.5440$).

[**B**] Spike–LFP coherence spectrograms for broad-spiking neurons.

[**C**] Mean spike–LFP coherence in the broadband gamma range (30–100 Hz) for broad-spiking neurons. LMM, $p = 0.2486$,

[**D**] Spike–LFP coherence spectrograms for narrow-spiking neurons.

[**F**] Mean spike–LFP coherence in the broadband gamma range (30–100 Hz) for narrow-spiking neurons LMM, $p = 0.6681$.

Statistical analyses were performed using LMM to account for variability across cells and animals. Bar plots show mean ± SEM. Dots represent individual neuron data. Number of female mice = 13. Total number of neurons (all): estrus = 178, estrus+Veh = 72. Broad spiking neurons: estrus = 110, estrus+Veh = 50. Narrow Spiking neurons: estrus= 68, estrus+Veh = 22.

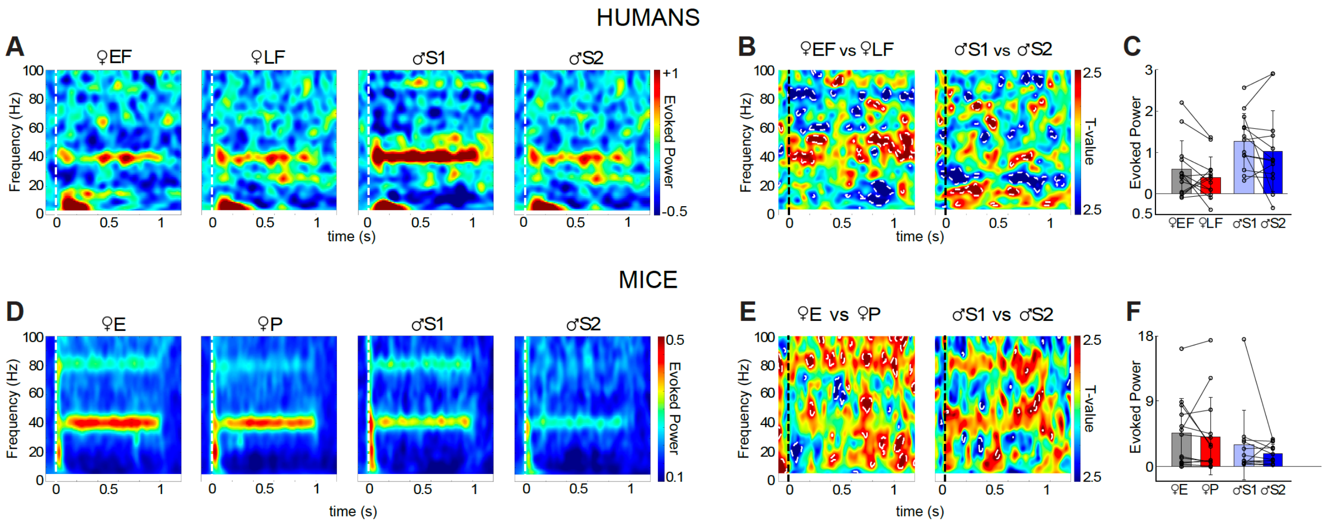


**Supplementary figure 6. Steady-state 40-Hz evoked power does not show estrous cycle modulation across species.**

[**A**] Baseline corrected, time–frequency spectrograms for each experimental group in humans.The onset of auditory stimulation is indicated by a white dotted line.

[**B**] Paired statistical maps showing differences in evoked power between conditions in humans Time–frequency regions exhibiting significant differences in power ($P < 0.05$) are outlined by white dotted lines; stimulus onset is indicated by a black dotted line.

[**C**] Evoked power at the stimulating frequency (40 Hz) in humans. Wilcoxon signed-rank: women EF vs LF: test $z=1.4327$, $p = 0.1520$; men S1 vs S2: $z=1.0832$, $p = 0.2787$.

[**D**] Baseline corrected, time–frequency spectrograms for each experimental group in naïve mice. The onset of auditory stimulation is indicated by a white dotted line.

[**E**] Paired statistical maps showing differences in evoked power between conditions in mice. Time–frequency regions exhibiting significant differences in power ($P < 0.05$) are outlined by white dotted lines; stimulus onset is indicated by a black dotted line.

[**F**] Evoked power at the stimulating frequency (40 Hz) in naïve mice. Wilcoxon signed-rank: females in estrus vs proestrus: $z = 0.5940$, $p = 0.5525$; males S1 vs S2: $z = 1.4905$, $p = 0.1361$.

In mice, only ASSR cooccurring with periods of immobility were analyzed. Bar plots show mean ± SEM. Dots represent individual data. EF: early follicular; LF: late follicular; E: estrous; P: proestrus. Number of human participants: women = 13, men = 13; number of mice: females = 13, males = 12.

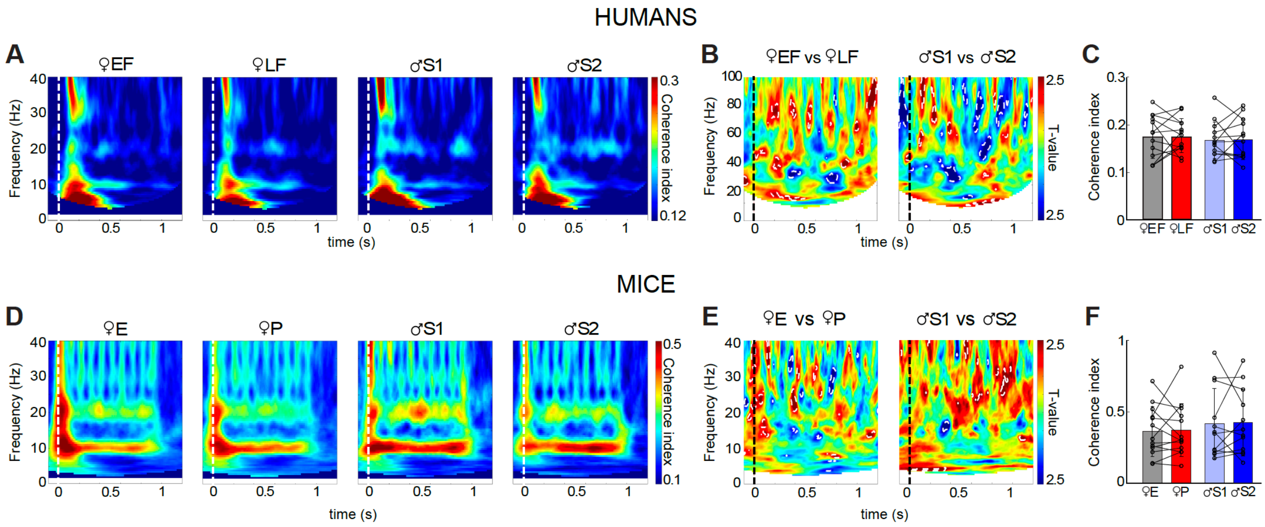


**Supplementary figure 7. Inter-trial coherence during steady state stimulation at 10Hz does not show estrous cycle modulation neither in humans nor in mice.**

[**A**] Time–frequency maps of inter-trial coherence for each experimental group in humans. The onset of auditory stimulation is indicated by a white dotted line.

[**B**] Paired statistical maps showing differences in ITC between conditions in humans. Time–frequency regions exhibiting significant differences ($P < 0.05$) are outlined by white dotted lines; stimulus onset is indicated by a black dotted line.

[**C**] Coherence index values corresponding to the stimulating frequency (10 Hz). Wilcoxon signed-rank: women EF vs LF: test $z=-0.2446$, $p = 0.8086$; men S1 vs S2: $z=-0.1048$, $p = 0.9165$.

[**D**] Time–frequency maps of inter-trial coherence for each experimental group in naïve mice. The onset of auditory stimulation is indicated by a white dotted line.

[**E**] Paired statistical maps showing differences in ITC between conditions in mice. Time–frequency regions exhibiting significant differences ($P < 0.05$) are outlined by white dotted lines; stimulus onset is indicated by a black dotted line.

[**F**] Coherence index values corresponding to the stimulating frequency (10 Hz). Wilcoxon signed-rank: females in estrus vs proestrus: $z = 0.1048$, $p = 0.9165$; males S1 vs S2: $z = -0.2353$, $p = 0. 8139$.

In mice, only ASSR cooccurring with periods of immobility were analyzed. Bar plots show mean ± SEM. Dots represent individual data. EF: early follicular; LF: late follicular; E: estrous; P: proestrus. Number of human participants: women = 13, men = 13; number of mice: females = 13; male = 12.

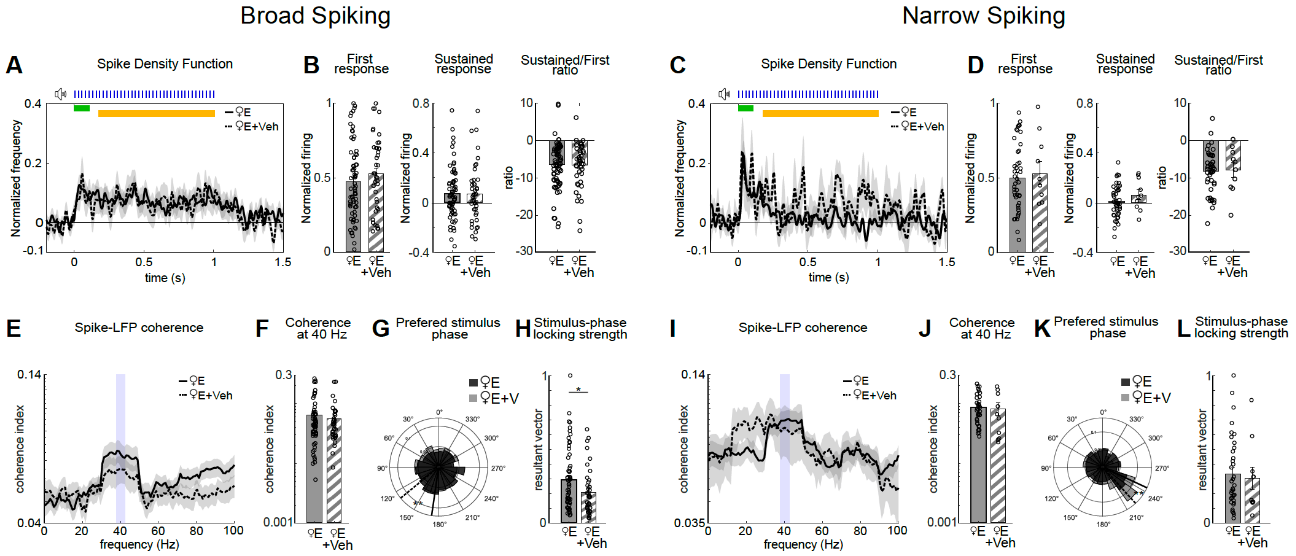


**Supplementary Figure 8. Vehicle injection at the estrus phase does not change firing response or spike-LFP coherence during 40 Hz auditory steady-state stimulation in mice.**

[**A**] Spike density functions (mean ± SEM, shaded) for responsive broad-spiking neurons in female mice during estrus and estrus+Vehicle. The early response window (0–100 ms post-stimulus) is indicated in green, and the sustained response window (200–1000 ms) in yellow.

[**B**] Firing rates of broad-spiking neurons during the early and sustained response windows, and their ratio (sustained/early). Acute vehicle injection had no effect on normalized firing during early response (LMM, p = 0.4898), sustained response (LMM, p=0.9636), or the sustained/early ratio (LMM, p=0.8224).

[**C**] Spike density functions (mean ± SEM, shaded) for responsive narrow-spiking neurons in female mice during estrus and estrus+Vehicle. The early response window (0–100 ms post-stimulus) is indicated in green, and the sustained response window (200–1000 ms) in yellow.

[**D**] Firing rates of narrow-spiking neurons during the early and sustained response windows, and their ratio (sustained/early). Acute vehicle injection had no effect on normalized firing during early response (LMM, p=0.5452), sustained response (LMM, p=0.6900), or their ratio (LMM, p=0.6396).

[**E**] Spike–LFP coherence spectra for broad-spiking responsive neurons.

[**F**] Spike–LFP coherence at 40 Hz for broad-spiking responsive neurons, LLM, p=0.9489.

[**G**] Circular histograms show different preferred firing phase at 40 Hz between groups for broad-spiking neurons (Watson-U2 test: U=0.2836, p=0.0074).

[**H**] Resultant vector in broad-spiking neurons (LLM, p=0.0397).

[**I**] Spike–LFP coherence spectra for narrow-spiking responsive neurons.

[**J**] Spike–LFP coherence at 40 Hz for narrow-spiking neurons. LLM, p = 0.8380.

[**K**] Circular histograms show different preferred firing phase at 40 Hz between groups for narrow-spiking neurons (Watson-U2 test: U=0.2914, p=0.0064).

[**L**] Resultant vector in narrow-spiking neurons (LLM, p=0.7064).

Bar plots show mean ± SEM. Dots represent individual neuron data. Broad Spiking cell number: estrus= 68, estrus + Veh= 58. Narrow Spiking cell number: estrus = 38, estrous+ Veh= 26. N female mice=13.

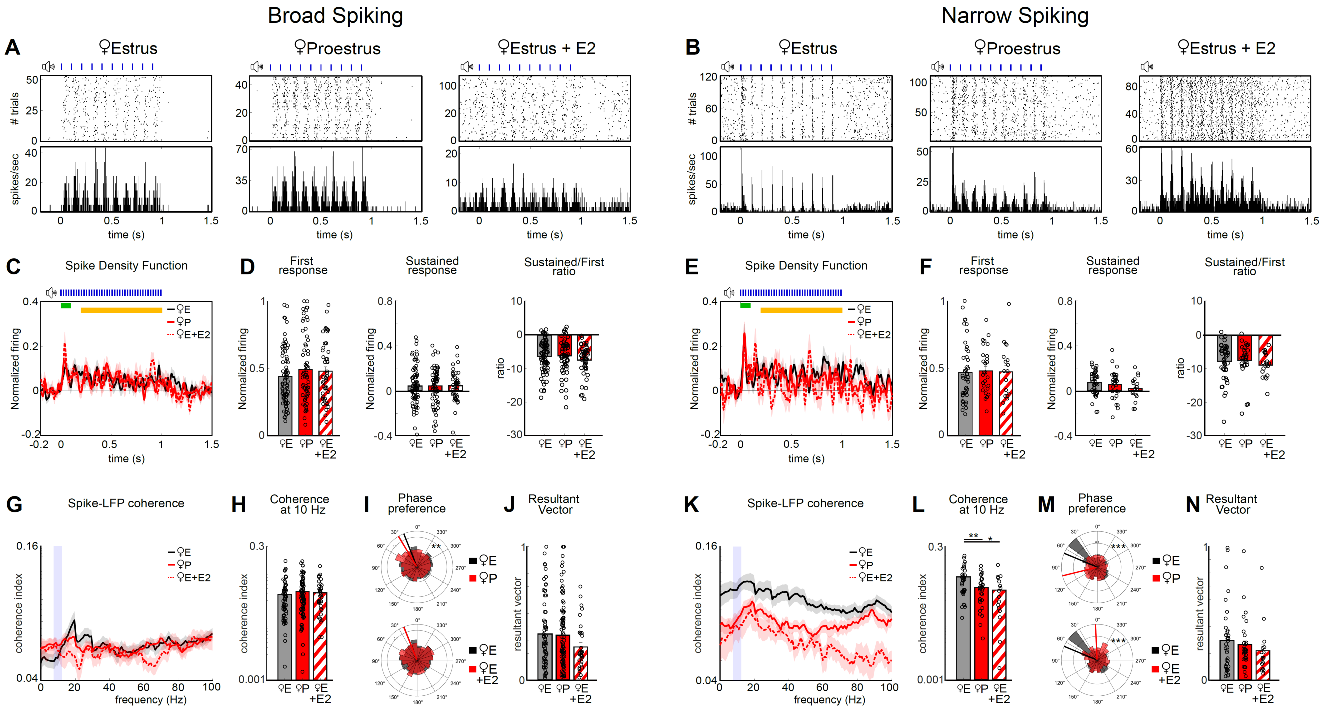


**Supplementary Figure 9. Estrous cycle modulates spike-LFP coherence of narrow band neurons during a 10 Hz auditory steady-state stimulation in mice.**

[**A**] Example raster plots (top) and peri-stimulus time histograms (bottom) for broad-spiking neurons during estrus (left), proestrus (middle) and estrus+E2 (right).

[**B**] Example raster plots (top) and peri-stimulus time histograms (bottom) for narrow-spiking neurons during estrus (left), proestrus (middle) and estrus+E2 (right).

[**C**] Spike density functions (mean ± SEM, shaded) for responsive broad-spiking neurons in female mice during estrus (black), proestrus (red) and estrus + E2 (dotted red). The early response window (0–100 ms post-stimulus) is indicated in green, and the sustained response window (200–1000 ms) in yellow.

[**D**] Firing rates of broad-spiking neurons during the early and sustained response windows, and their ratio (sustained/early). Normalized firing shows no differences during early response (estrous vs proestrus: LMM, $p = 0.2092$; estrus vs estrus+E2: LLM, $p=0.3092$), sustained response (estrous vs proestrus: LMM, $p = 0.8984$; estrus vs estrus+E2: LLM, $p=0.9365$), or the sustained/early ratio (estrous vs proestrus: LMM, $p = 0.9089$; estrus vs estrus+E2: LLM, $p=0.2948$).

[**E**] Spike density functions (mean ± SEM, shaded) for responsive narrow-spiking neurons in female mice during estrus (black), proestrus (red) and estrus + E2 (dotted red). The early response window (0–100 ms post-stimulus) is indicated in green, and the sustained response window (200–1000 ms) in yellow.

[**F**] Firing rates of narrow-spiking neurons during the early and sustained response windows, and their ratio (sustained/early). Normalized firing show no differences between groups during early response (estrous vs proestrus: LMM, p = 0.7921; estrus vs estrus+E2: LLM, p=0.6474), sustained response (estrous vs proestrus: LMM, p = 0.9980; estrus vs estrus+E2: LLM, p=0.0725), ratio (estrous vs proestrus: LMM, p = 0.8382; estrus vs estrus+E2: LLM, p=0.6134)

[**G**] Spike–LFP coherence spectra for broad-spiking responsive neurons.

[**H**] Spike–LFP coherence at 10 Hz of broad-spiking neurons (estrous vs proestrus: LMM, p = 0.1304; estrus vs estrus+E2: LLM, p=0.6706)

[**I**] Circular histograms show different preferred firing phase at 10 Hz for broad-spiking neurons (Watson-U2 test, estrus vs proestrus: U=0.3075, p=0.0046; estrus vs estrus+E2: U=0.0424, p=0.8659).

[**J**] Resultant vector in broad-spiking neurons (LLM, estrus vs proestrus: p=0.8897; estrus vs estrus+E2: p=0.0927).

[**K**] Spike–LFP coherence spectra for narrow-spiking responsive neurons.

[**L**] Spike–LFP coherence at 10 Hz of narrow-spiking neurons shows more spike-LFP coherence during estrus than proestrus (LMM, p = 0.0089) and estrous+E2 (LLM, p=0.0277).

[**M**] Circular histograms show different preferred firing phase at 10 Hz for narrow-spiking neurons (Watson-U2 test, estrus vs proestrus: U=0. 4408, p=3.3251*10-4; estrus vs estrus+E2: U=0.4345, p=3.7675*10-4).

[**N**] Resultant vector in narrow-spiking neurons (LLM, estrus vs proestrus: p=0.6555; estrus vs estrus+E2: p=0.5798).

*P<0.05, **P<0.01. Bar plots show mean ± SEM. Dots represent individual neuron data. Broad Spiking cell number: estrus= 70, proestrus= 90, estrus+E2= 37. Narrow Spiking cell number: estrus= 36, proestrus= 33, estrous+E2= 16. N female mice=13.
